# DpaA detaches Braun’s lipoprotein from peptidoglycan

**DOI:** 10.1101/2021.02.21.432140

**Authors:** Matthias Winkle, Víctor M. Hernández-Rocamora, Karthik Pullela, Emily C. A. Goodall, Alessandra M. Martorana, Joe Gray, Ian R. Henderson, Alessandra Polissi, Waldemar Vollmer

## Abstract

Gram-negative bacteria have a unique cell envelope with a lipopolysaccharide-containing outer membrane that is tightly connected to a thin layer of peptidoglycan. The tight connection between the outer membrane and peptidoglycan is needed to maintain the outer membrane as an impermeable barrier for many toxic molecules and antibiotics. Enterobacteriaceae such as *Escherichia coli* covalently attach the abundant outer membrane-anchored lipoprotein Lpp (Braun’s lipoprotein) to tripeptides in peptidoglycan, mediated by the transpeptidases LdtA, LdtB and LdtC. LdtD and LdtE are members of the same family of LD-transpeptidases but they catalyse a different reaction, the formation of 3-3 cross-links in the peptidoglycan. The function of the sixth homologue in *E. coli*, LdtF remains unclear, although it has been shown to become essential in cells with inhibited LPS export to the outer membrane. We now show that LdtF hydrolyses the Lpp-peptidoglycan linkage, detaching Lpp from peptidoglycan, and have renamed LdtF to peptidoglycan *meso*-**d**iamino**p**imelic **a**cid protein **a**midase A (DpaA). We show that the detachment of Lpp from peptidoglycan is beneficial for the cell under certain stress conditions and that the deletion of *dpaA* allows frequent transposon inactivation in the *lapB (yciM)* gene, whose product down-regulates lipopolysaccharide biosynthesis. DpaA-like proteins have characteristic sequence motifs and are present in many Gram-negative bacteria of which some have no Lpp, raising the possibility that DpaA has other substrates in these species. Overall, our data show that the Lpp-peptidoglycan linkage in *E. coli* is more dynamic than previously appreciated.

**IMPORTANCE:** Gram-negative bacteria have a complex cell envelope with two membranes and a periplasm containing the peptidoglycan layer. The outer membrane is firmly connected to the peptidoglycan by highly abundant proteins. The outer membrane-anchored Braun’s lipoprotein (Lpp) is the most abundant protein in *E. coli* and about one third of the Lpp molecules become covalently attached to tripeptides in peptidoglycan. The attachment of Lpp to peptidoglycan stabilizes the cell envelope and is crucial for the outer membrane to function as a permeability barrier for a range of toxic molecules and antibiotics. So far the attachment of Lpp to peptidoglycan has been considered to be irreversible. We have now identified an amidase, DpaA, which is capable of detaching Lpp from PG and we show that the detachment of Lpp is important under certain stress conditions. DpaA-like proteins are present in many Gram-negative bacteria and may have different substrates in these species.

## INTRODUCTION

Bacteria have to maintain the integrity of their complex cell envelope at all times when propagating in diverse and often adverse environments (1). The peptidoglycan (PG) layer surrounds the cytoplasmic membrane (CM) and protects the cell from rupture due to osmotic challenges (2, 3). Diderm (Gram-negative) bacteria protect themselves from antimicrobial compounds by surrounding their PG layer by a semi-permeable outer membrane (OM). In *Escherichia coli* and related species, the PG and OM are tightly connected by multiple, highly abundant proteins such as Lpp (Braun’s lipoprotein), OmpA and Pal (4–7). The cell must maintain these tight connections at all times and coordinate the expansion of the PG and OM during growth and cell division to avoid leaks in the OM, the loss of OM vesicles or lysis due to an instable cell envelope (6, 8, 9).

Lpp is the most abundant protein in *E. coli* with about 1 million copies per cell (5). In pioneering work in the 1970s, Volkmar Braun and co-workers identified Lpp not only as the first protein that is covalently attached to PG, but also as the first protein with a lipid modification (4). The Lpp pre-protein uses the Sec system for translocation through the CM where it then matures by (i) the addition of a diacylglycerol to a cysteine residue near the N-terminus, (ii) the removal of the leader peptide and (iii) the addition of a third acyl chain to the N-terminus (10). Mature Lpp is then transported through the periplasm and anchored to the inner leaflet of the OM by the localisation of lipoprotein (Lol) transport system (11). About one third of the protein molecules are covalently attached to PG via the ε-amino group of the C-terminal lysine, which is linked to the α-carboxylic group of *meso*-diaminopilemic acid (*m*DAP) at position 3 of a PG stem peptide, by the LD-transpeptidases (LDTs) LdtA, LdtB and LdtC (4, 12, 13). About 5% of all PG subunits are attached to Lpp in exponentially growing cells and 15% in stationary phase (14).

In addition to stabilizing the cell envelope, Lpp helps to maintain a constant distance between the PG and OM (15, 16). Mutants lacking Lpp or LdtA, LdtB and LdtC suffer from hyper-vesiculation, losing OM-derived vesicles filled with periplasmic content into the environment (9, 17, 18). These mutants become sensitive to sodium dodecyl sulphate (SDS) due to abnormally high OM permeability (6). Pathogenic *E. coli* and *Salmonella* mutants lacking Lpp are less virulent and able to invade their host, and an Lpp-deficient uropathogenic *E. coli* strain is more susceptible to serum killing (19–21). In this strain, the attachment of Lpp to PG was crucial for the expression of capsular polysaccharide, illustrating another key role of PG-attached Lpp (21).

Given the important role of Lpp for cell envelope stability in *E. coli*, it was surprising that Lpp is only present in closely related γ-proteobacteria (22). While another PG-attached lipoprotein has been described in *Pseudomonas* spp., the covalent attachment of PG to OM localized proteins has not been studied in depth in other diderm bacteria PG (23). This changed recently when two publications reported the covalent attachment of certain OM β-barrel proteins (OMPs) in a wide range of α-proteobacteria (24, 25). Additionally to OMPs, one of the studies identified the OM-anchored lipoprotein LimB attached to PG in *Coxiella burnetti* (25). Similar to Lpp in *E. coli*, these proteins are attached to the *m*DAP residue in PG stem peptides by LDTs and are required for OM stability. Different to Lpp, OMPs were found to be attached to PG via N-terminal glycine or alanine residues. Many Firmicutes (Gram-positives) attach multiple cell wall proteins to PG using the enzyme sortase (26). These are often involved in interactions with host factors during bacterial infections and include, for example, protein A in *S. aureus* or pilus subunits in *S. pneumoniae* (27, 28).

Although previous work identified several PG-attached proteins and two classes of enzymes for protein-PG attachment (LDTs and sortase), to date no enzyme for the opposite reaction, the detachment of proteins from PG, has been identified. Here, we show that an LDT homologue of previously unknown function (DpaA, formerly named LdtF or YafK), has an amidase activity hydrolysing the bond between Lpp and PG in *E. coli*. We also report that the DpaA-mediated detachment of Lpp from PG helps the cell to cope with certain stress conditions and that DpaA-like enzymes are present in many diderm bacteria.

## RESULTS

### *DpaA* is genetically linked to PG and OM biogenesis

In *E. coli*, all six LD-transpeptidases (LDTs) that contain the YkuD conserved motif (**Fig. 1**) become important under cell envelope stress conditions. LdtA, LdtB and LdtC facilitate cell envelope stability by covalently attaching the OM-anchored lipoprotein Lpp to PG (13). LdtD forms 3-3 cross-links in PG to repair the PG layer upon defective OM assembly (29). LdtE appears to form 3-3 cross-links when cells enter the stationary phase of growth, although its activity has not been demonstrated with the purified enzyme. The function of the last LDT, LdtF (now DpaA), remained unknown. Genetic depletion of the essential LPS export gene *lptC* causes lysis in cells lacking *dpaA* (29), but these show morphological defects even in the presence of inducer (arabinose), and the lysis of *lptC*-depleted Δ*dpaA* cells can be prevented by the additional deletion of a novel amidase activator gene, *actS* (Gurnani *et al.,* under review). None of these phenotypes are observed for the other *ldt* mutants, suggesting that DpaA has different (or additional) roles than LdtA-E.

**Figure 1.**
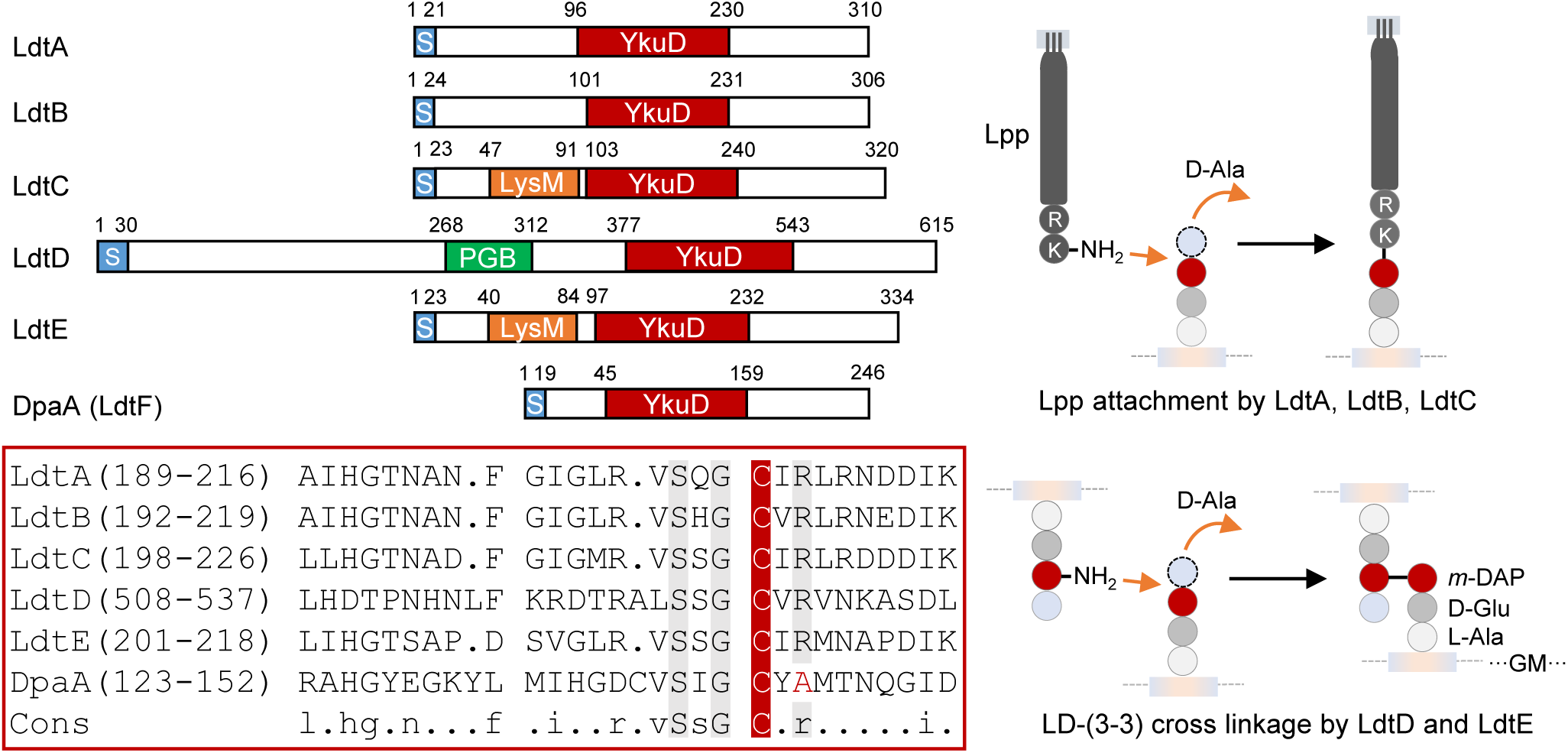
Overview of the different LDT family members. *E. coli* has six proteins with a YkuD (LDT) domain (top left). Protein regions: S, signal peptide (S); LysM and PGB, peptidoglycan-binding domains. The numbers indicate amino acid positions. Sequence alignment within the YkuD domain (bottom left) shows the active site cysteine (C, in red) and other conserved amino acid residues in grey. DpaA (LdtF) misses a conserved arginine (R) and its function was unknown. Right side: LdtA, LdtB and LdtC attach Lpp to *m*DAP in PG stem peptides. LdtD and presumably LdtE form 3-3 cross-links in PG.

To better understand the role of DpaA we first performed genome-wide transposon insertion analysis (TraDIS) of the *dpaA* mutant. We constructed two libraries of transposon mutants by electroporation of a mini-Tn5 transposon into a Δ*dpaA* strain, and the BW25113 parent strain as a control. To identify the transposon insertion sites, we sequenced the transposon-genomic DNA junction and mapped the data to the BW25113 reference genome as previously described (62). We identified 670,433 and 360,972 unique insertion sites for *dpaA* and control library, respectively, and these are distributed around the chromosome (**Fig. S1**, **Table S1**). There were no insertions within *dpaA* in the Δ*dpaA* library, confirming the library genotype (**Fig. 2B**). To identify mutants with a fitness defect, or advantage, in the *dpaA* gene-deletion background, we used the BioTraDIS EdgeR analysis tool to compare the frequency of reads per gene between each library, using thresholds of >2-fold change and Q-value > 0.01. This analysis revealed relatively few genes with significantly more or less reads in the *dpaA* mutant compared to wild-type. The most obvious differences we observed in the *dpaA* library compared to the control were a 4.25 fold enrichment (Q-value = 1.84E-25) of reads within *lapB* (*yciM*) and a 2.1 fold enrichment of reads with the neighbouring *lapA* (yciS) gene (Q-value = 2.59E-5) (**Fig. 2A and 2C**). LapA and LapB are essential regulators of LPS biosynthesis, regulating the degradation of LpxC by the membrane-bound protease FtsH (30, 31) and LapA physically interacts with the elongasome (31b). Hence, *dpaA* inactivation is linked to phenotypes related to both LPS synthesis (*lapB* inactivation), peptidoglycan synthesis (*lapA* inactivation) and LPS export (*lptC* depletion). To begin to understand these genetic interactions, we wanted to determine the enzymatic activity of DpaA.

**Figure 2.**
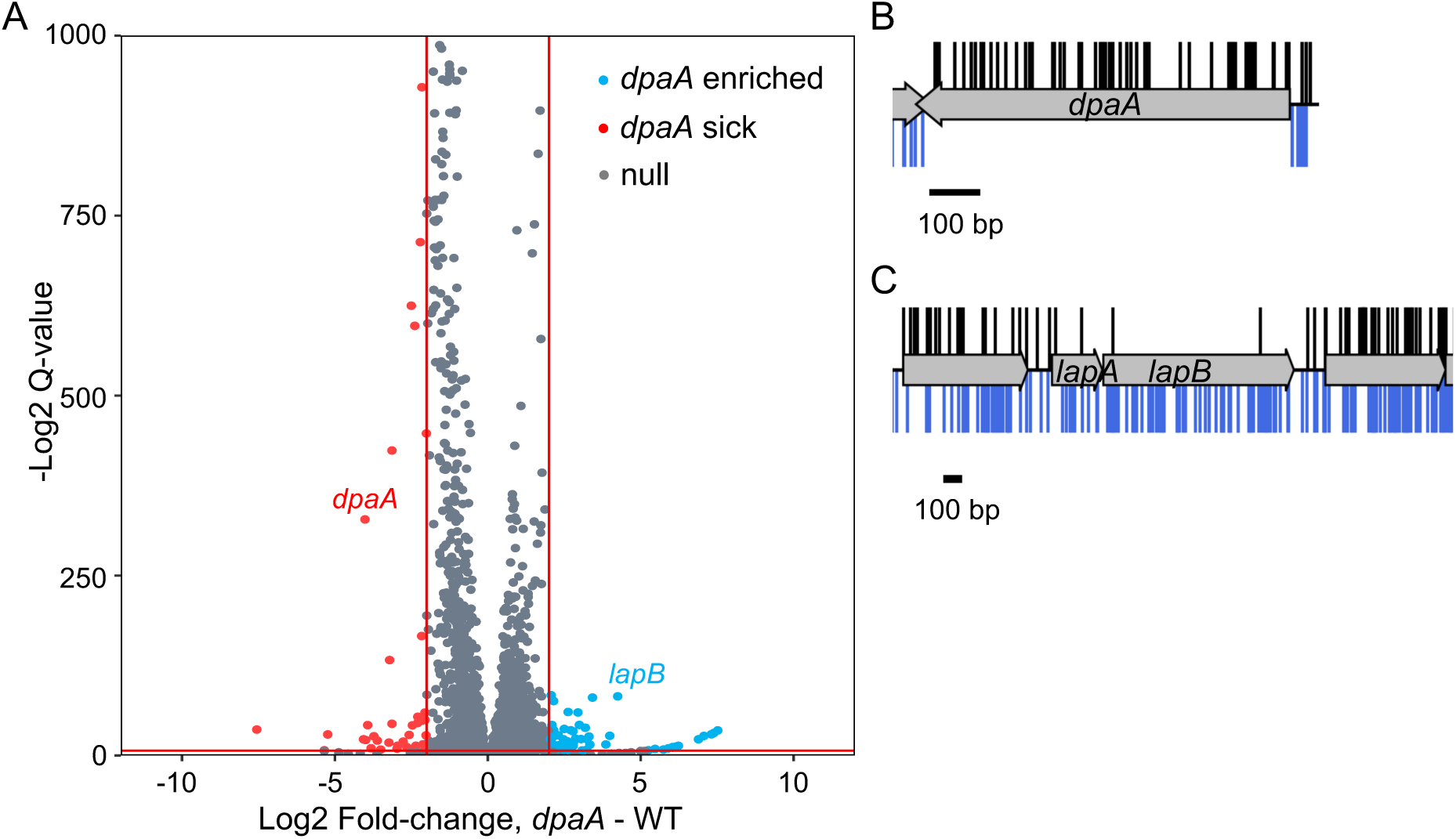
Comparison of the read depth between genes in each transposon library. (**A**) Volcano plot showing a comparison of the relative mutant abundance between libraries. Genes with a >2-fold change in read abundance and with a Q-value > 0.01 are coloured; these thresholds are represented as solid red lines. (**B**,**C**) The insertion profile of *dpaA* (B) and the *lapAB* operon (C) for both libraries, with transposon insertion sites identified in the control library displayed above the gene track (black) and transposon insertion events in the *dpaA* library shown below the gene track (blue). The insertion plot is capped at a read frequency of 1, scale bar = 100 bp.

### DpaA is not an LD-TPase

We purified LdtD, LdtE and DpaA and first tested for their ability to produce 3-3 cross-links in PG isolated from an *E. coli* strain lacking all 6 LDTs (BW25113Δ6LDT) at neutral (pH 7.5) or acidic (pH 5.0) conditions. The PG was digested with the muramidase cellosyl and the resulting muropeptides were separated by high performance liquid chromatography (HPLC) (**Fig. S2A and S2B**). LdtD produced muropeptides with 3-3 cross-links or tripeptides [TetraTri(3-3), TriTri(3-3), TetraTri, Tri; Fig. S2C)], confirming LD-TPase and LD-CPase activities (29). LdtE generated TetraTri(3-3) consistent with the presence of this TPase products in the PG of BW25113Δ6LDT cells overexpressing *ldtE* and *dpaA* (32). LdtE was nearly inactive at pH 7.5 and did not show LD-CPase acitvity. DpaA did not show any LD-TPase or LD-CPase activity under any condition tested, despite long incubation times of 12 hours. We noticed that DpaA lacked an active-site arginine residue conserved in other LDTs (**Fig. 1**), suggesting that DpaA is an inactive enzyme of the LDT family or has a different activity than the other LDTs.

### DpaA releases Lpp from PG

We reasoned that rather than forming LD-bonds, DpaA might hydrolyse bonds generated by other LDTs, however, in several experiments we could not detect any LD-endopeptidase or LD-carboxypeptidase activity (not shown). Hence, we next tested if DpaA hydrolyses the bond between the terminal L-lysine residue of Lpp and PG, which is formed by LdtA-C (13). We first used purified muropeptides containing the terminal two amino acids (lysine and arginine) from Lpp, as artificial substrates. The protease pronase E used in PG purification removes contaminating proteins and most part of the covalently attached Lpp, with the exception of the C-terminal Lys-Arg dipeptide; the residual dipeptide is used in PG analysis to quantify the amount of PG-attached Lpp in strains (14).

We incubated Tri-LysArg and TetraTri-LysArg with DpaA or the catalytically inactive DpaA(C143A) (with Ala replacing the active site Cys residue) and analysed the products by HPLC. DpaA but not DpaA(C143A) converted both substrates to muropeptides lacking the Lys-Arg dipeptide (Tri or TetraTri) (**Fig. 3A** **and S2D**). We confirmed the mass of the substrate and product muropeptides by MS/MS analysis (**Fig. S3**). To test if this activity was specific for DpaA we incubated Tri-LysArg with purified LdtD or LdtE and observed no activity with these LDTs (**Fig. S2E**).

**Figure 3.**
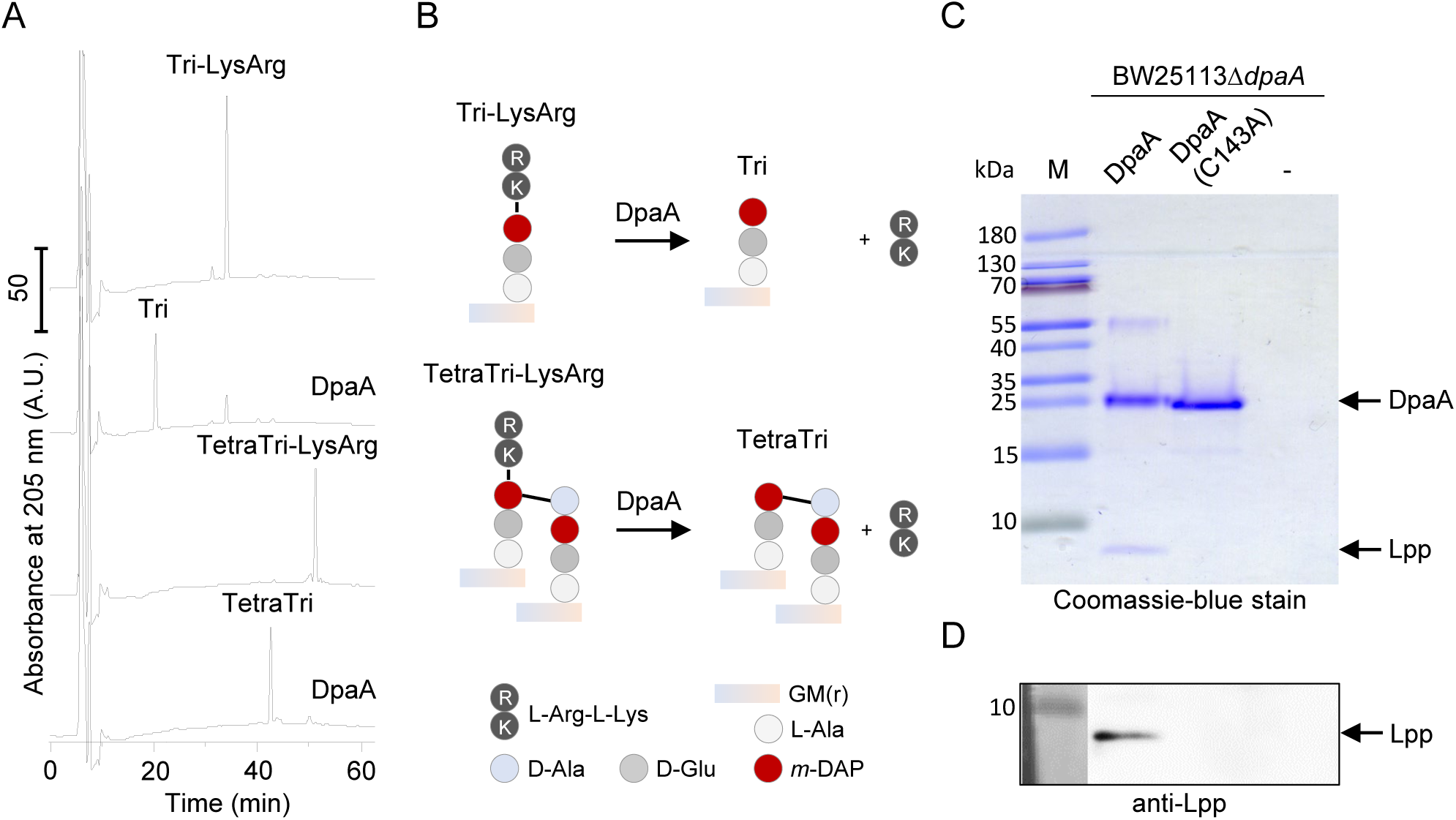
DpaA detaches Lpp from PG. (**A**) Activity against the muropeptides Tri-LysArg and TetraTri-LysArg. Muropeptides were incubated with DpaA and the reaction products reduced with sodium borohydride and separated by HPLC. DpaA hydrolysed both substrates, releasing the muropeptides lacking L-Arg-L-Lys. The released dipeptide is missing in the chromatogram because it presumably co-elutes early with the salts at ∼8 min. Fig. S3 shows the MS data proving the identities of the muropeptide substrates used here, and of the products. (**B**) Scheme of the reactions observed in A. Data in Fig. S2E show that LdtD and LdtE are not active against Tri-LysArg. Fig. S4A,B shows the inhibition of DpaA by copper. (**C, D**) Lpp release assay. DpaA was incubated with PG sacculi from BW25113*ΔdpaA* that were not treated with pronase E and hence contained PG-attached Lpp. Reaction mixtures were boiled and proteins were separated by SDS-PAGE, following by staining with Coomassie-blue (**C**) or Western Blotting and detection of Lpp with specific antiserum (**D**). DpaA, but not the catalytically inactive DpaA(C143A), released Lpp from PG. M, protein size marker.

LDTs are known to be inhibited by copper ions (33). To test if this was true for DpaA, we incubated PG from BW25113*ΔdpaA* cells with DpaA in the presence of CuCl_2_, followed by the quantification of Lys-Arg containing muropeptides by HPLC. In the absence of copper, DpaA removed Tri-LysArg from the muropeptide profile, forming Tri whilst the presence of 0.2 mM CuCl_2_ inhibited the reaction (**Fig. S4A**). We observed half-inhibition at ∼0.1 mM CuCl_2_ (**Fig. S4B**), indicating an approx. 6-fold lower IC_50_ than that for LdtD (33).

In a different assay for DpaA activity, we incubated the protein with PG sacculi from BW25113*ΔdpaA* cells that were not treated with pronase E and, hence contained covalently attached full-length Lpp (rather than only the Lys-Arg dipeptide). Samples were centrifuged to separate soluble proteins from insoluble PG, and proteins present in the supernatant were analysed by SDS-PAGE (**Fig. 3C**) and Western Blot developed with a specific antibody against Lpp (**Fig. 3D**). DpaA, but not inactive DpaA(C143A) was capable of releasing Lpp from PG, confirming the results obtained with the soluble muropeptides with Lys-Arg dipeptides. Altogether, our data prove that DpaA hydrolyses the bond between *m*DAP in PG and the C-terminal Lys residue of Lpp, releasing Lpp from the PG.

### DpaA has also CPase activity against glycine-containing muropeptides

When we further investigated the activity of DpaA. We found that it released LysArg dipeptides from PG under acidic or neutral conditions (**Fig. 4A**). In the course of these experiments we noticed that DpaA also acted on rare peptides in PG with glycine (instead of D-Ala) at position 4, for example TetraGly4 (Peak 2) or its dimer version TetraTetraGly4 (Peak 5) (14). These structures were hydrolysed by DpaA particularly at acidic conditions via a *m*DAP-Gly carboxypeptidase (CPase) activity.

**Figure 4.**
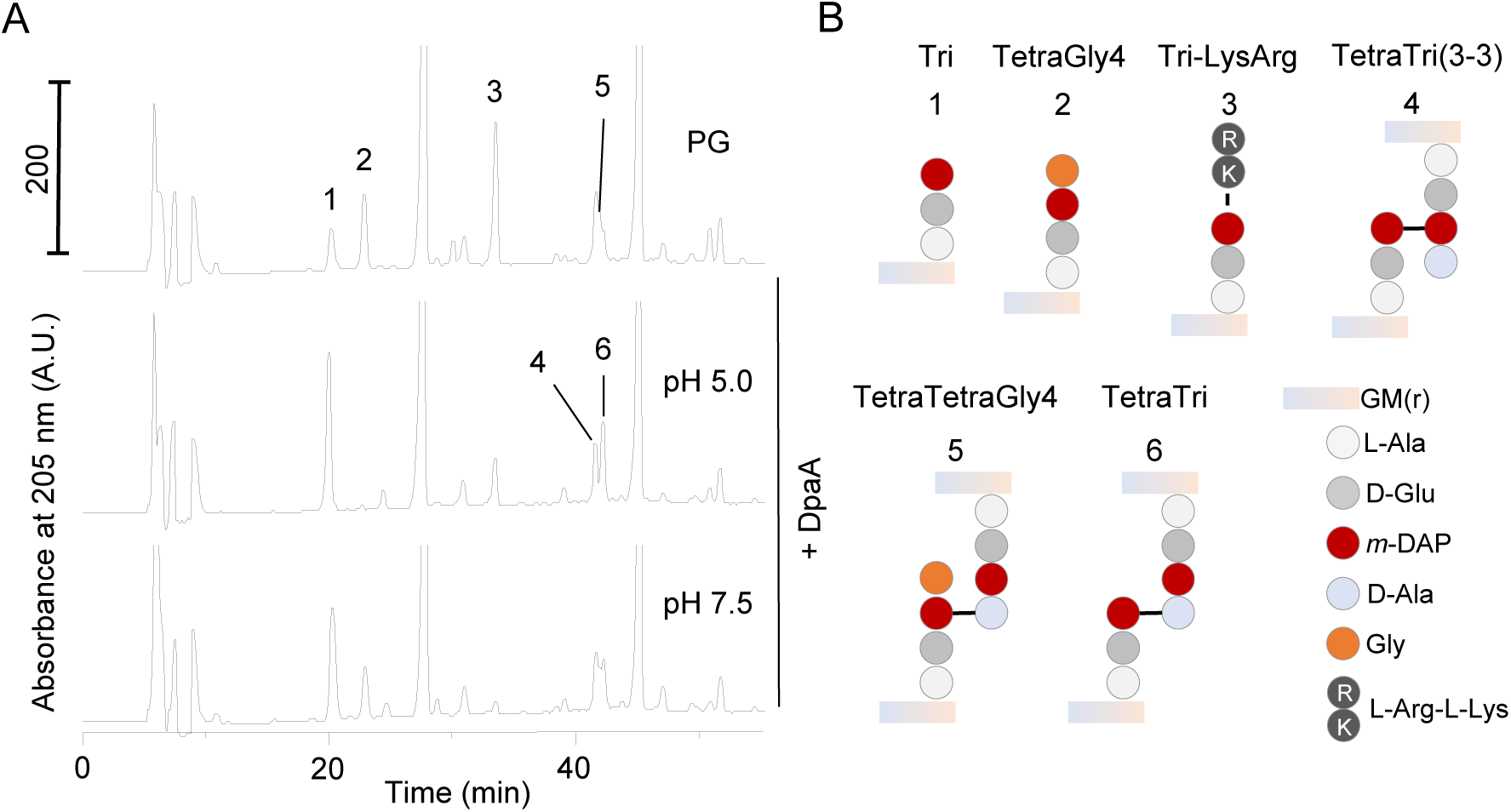
DpaA is active against TetraGly4 and TetraTetraGly4. (**A**) PG from strain *araB*p*lptC* Δ*dpaA* (+arabinose) was incubated with DpaA at pH 5.0 or 7.5, followed by digestion with cellosyl and analysis of the muropeptide composition by HPLC. In addition to its expected decrease of Tri-LysArg, DpaA was also active against muropeptides containing glycine at position 4 (muropeptides 2 and 5). DpaA showed higher activity at acidic pH. Chromatographs were cropped at 250 mAU to better observe the minor muropeptides. (**B**) Structures of the relevant muropeptide labelled in A. Fig. S4C shows that DpaA is not active against muropeptides with non-canonical D-amino acids at position 4 of the PG stem peptide.

We next tested if DpaA could remove other unusual amino acids present at the terminus of a tetrapeptide. For this, we used LdtD to exchange the terminal D-alanine residue by D-lysine, D-glutamine, D-valine or glycine in PG from BW25113Δ6LDT and tested if these can be removed by DpaA (**Fig. S4C**). As before, DpaA was active against muropeptides with glycine but not against those with D-lysine, D-glutamine or D-valine. Together, these data show that DpaA specifically removes Lys or Gly residues from position 4 of tetrapeptides, but not D-alanine or any of the other D-amino acids tested.

### DpaA is active in cells

Our biochemical experiments provided strong evidence that DpaA is able to release Lpp from PG. To test if this reaction can happen in cells, we ectopically expressed plasmid-borne *dpaA* or *dpaA(C143A)* in strains with deletions of the *dpaA*, *lpp* or *ldtABC* genes, and analysed the PG composition of exponentially growing cells (**Fig. 5A and 5B**). Expression of DpaA in BW25113*ΔdpaA* resulted in a two-fold reduction of Tri-LysArg from 3.7 ± 0.1% to 1.6 ± 0.0%, compared to BW25113 expressing the empty plasmid (pGS100) (**Fig. 5A and 5C**). BW25113*ΔdpaA* expressing catalytically inactive DpaA(C143A) had an increased level of Tri-LysArg compared to BW25113 pGS100, possibly due to the lack of DpaA activity. Tripeptides were increased in all samples with decreased LysArg-muropeptides due to the expression of active DpaA, for example, BW25113*ΔdpaA* pDpaA contained 11.0 ± 0.3% tripeptides compared to 4.9 ± 0.4% in BW25113 pGS100. This result suggests that cells counteract the high DpaA activity by increasing the activities of LdtA-C to maintain a sufficient amount of PG-attached Lpp.

**Figure 5.**
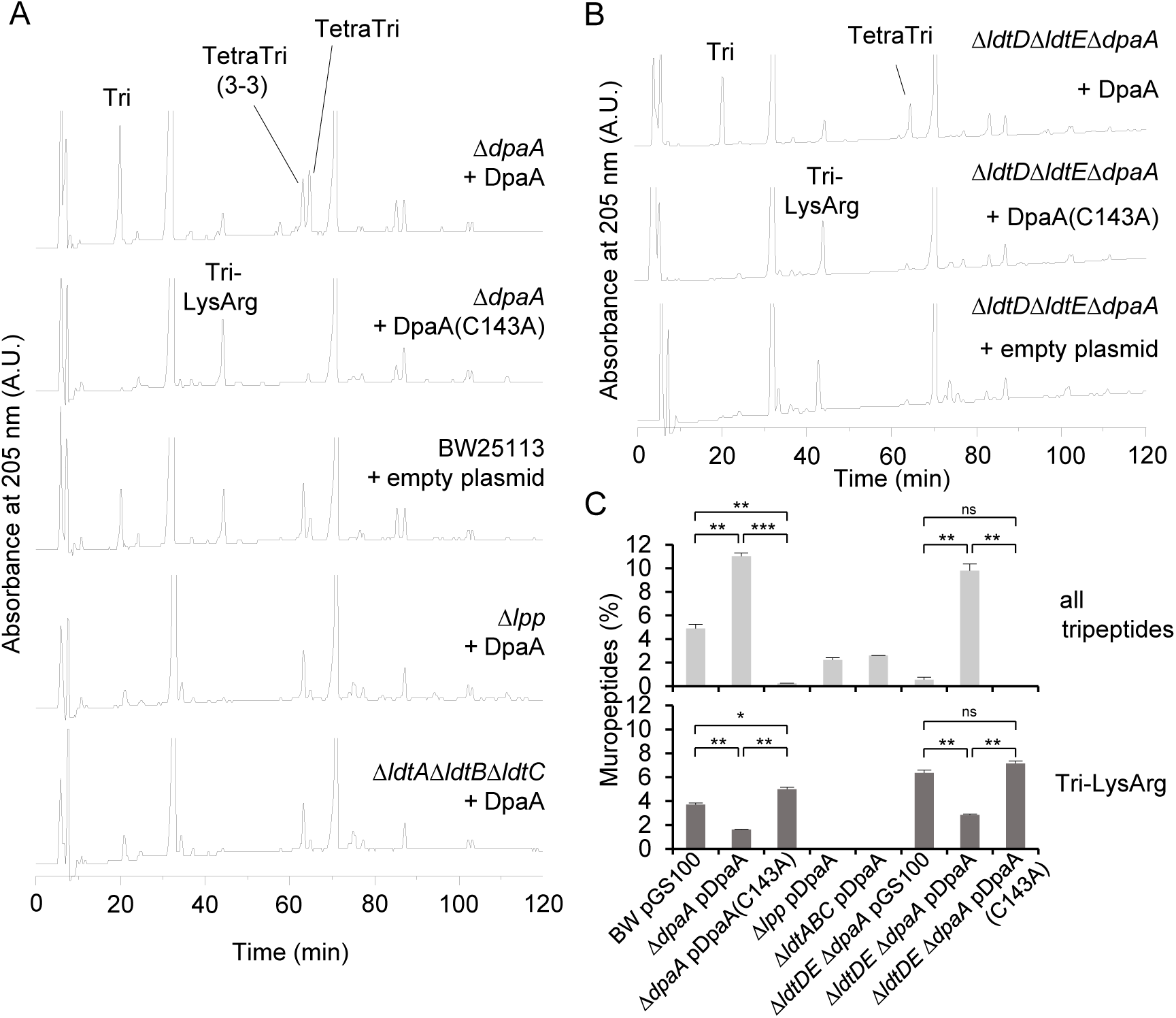
DpaA is active in cells. (**A**) Wild-type (BW25113) or mutants with single or multiple gene deletions (BW25113Δ*dpaA*, BW25113Δ*lpp*, BW25113Δ*ldtA*Δ*ldtB*Δ*ldtC*) expressing DpaA or DpaA(C143A) from a plasmid, or carrying the empty plasmid, were grown in LB medium, harvested, and the muropeptide composition was determined. Only expression of active DpaA, but not the inactive DpaA(C143A), reduced the Tri-LysArg peak when present. (**B**) Muropeptide profiles of BW25113Δ*ldtD*Δ*ldtE*Δ*dpaA* (lacking LDTs involved with 3-3 cross-link formation and DpaA) expressing DpaA, DpaA(C143A) or no DpaA version. Chromatographs were cropped above 250 mAU, the uncropped HPLC chromatographs of A and B are shown in Fig. S5. (**C**) Quantification of the total tripeptides and Tri-LysArg in the chromatograms shown in A and B. The values are mean ± variation in from two independent biological replicates, except BW25113Δ*ldtA*Δ*ldtB*Δ*ldtC* pDpaA, which was prepared once. Significance was measured by a two-tailed, homoscedastic T-test. ns (not significant), p > 0.05; *, p ≤ 0.05; **, p ≤ 0.01; ***, p ≤ 0.001.

The expression of DpaA had no significant effect on the PG composition in cells lacking Lpp (BW25113*Δlpp*) or the Lpp-attaching enzymes (BW25113*ΔldtAΔldtBΔldtC*) which, as expected, did not show detectable level of Tri-LysArg and therefore lack the substrate of DpaA (**Fig. 5A**). To test if LdtD or LdtE affect the activity of DpaA, we also analysed the PG composition of BW25113*ΔldtDΔldtEΔdpaA* cells containing pDpaA, pDpaA(C143A) or pGS100 (**Fig. 5B**). As expected, the PG of these cells did not contain 3-3 cross-links. The level of Tri-LysArg was higher in strains lacking *ldtD* and *ldtE*, presumably because LdtA-C compete with LdtD-E for tetrapeptide donors in PG. However, the expression of active DpaA correlated with a reduction in Tri-LysArg and an increase in tripeptides in PG from BW25113 and BW25113*ΔldtDΔldtEΔdpaA*, showing that the absence of LdtD and LdtE does not affect DpaA. Altogether these data verify that DpaA detaches Lpp from PG in cells.

### DpaA contributes to mecillinam resistance

Our previous work showed that DpaA is important in cells that experience severe OM assembly stress. Here we wanted to further explore conditions at which DpaA becomes important and searched a chemical genomics database for conditions that decreased the fitness of the *dpaA* mutant strain (34). The *dpaA* mutant had negative fitness scores with the inhibitor of LPS synthesis, CHIR-090, and the elongasome inhibitors A22 and mecillinam which both cause cells to become spherical (35, 36). We hypothesised that the tight connection between the OM and PG via PG-attached Lpp could complicate changes in cell shape such as the transition of rod-shaped to spherical shape. To test this hypothesis, we measured the sensitivity of strains towards mecillinam. When comparing the growth of BW25113, BW25113*ΔdpaA,* BW25113*Δlpp,* and BW25113*ΔdpaAΔlpp* on LB agar plates supplemented with mecillinam we observed a higher susceptibility of the *ΔdpaA* mutant (**Fig. 6A**). We also grew these strains in liquid LB medium in the presence or absence of 1 µg/ml mecillinam. Cells of BW25113*Δlpp* still grew 8.5 h post addition of mecillinam, while BW25113*ΔdpaAΔlpp* cells arrested growth, and the optical density of BW25113 and BW25113*ΔdpaA* cultures declined (**Fig. 6B**). Expressing *ftsQAZ* from pTB63 decreased the susceptibility to mecillinam as described (37), but the difference between the strains persisted. BW25113Δ*dpaA* pTB63 was more susceptible to mecillinam than the other strains and expressing pDpaA form plasmid restored mecillinam resistance (**Fig. S6B**). This effect was specific to mecillinam and the Δ*dpaA* strain showed similar susceptibility to tetracycline and ampicillin, which do not cause cells to become spherical (**Fig. S6A**). Hence, detachment of Lpp from PG appears to be beneficial for cells that undergo radical cell shape changes.

**Figure 6.**
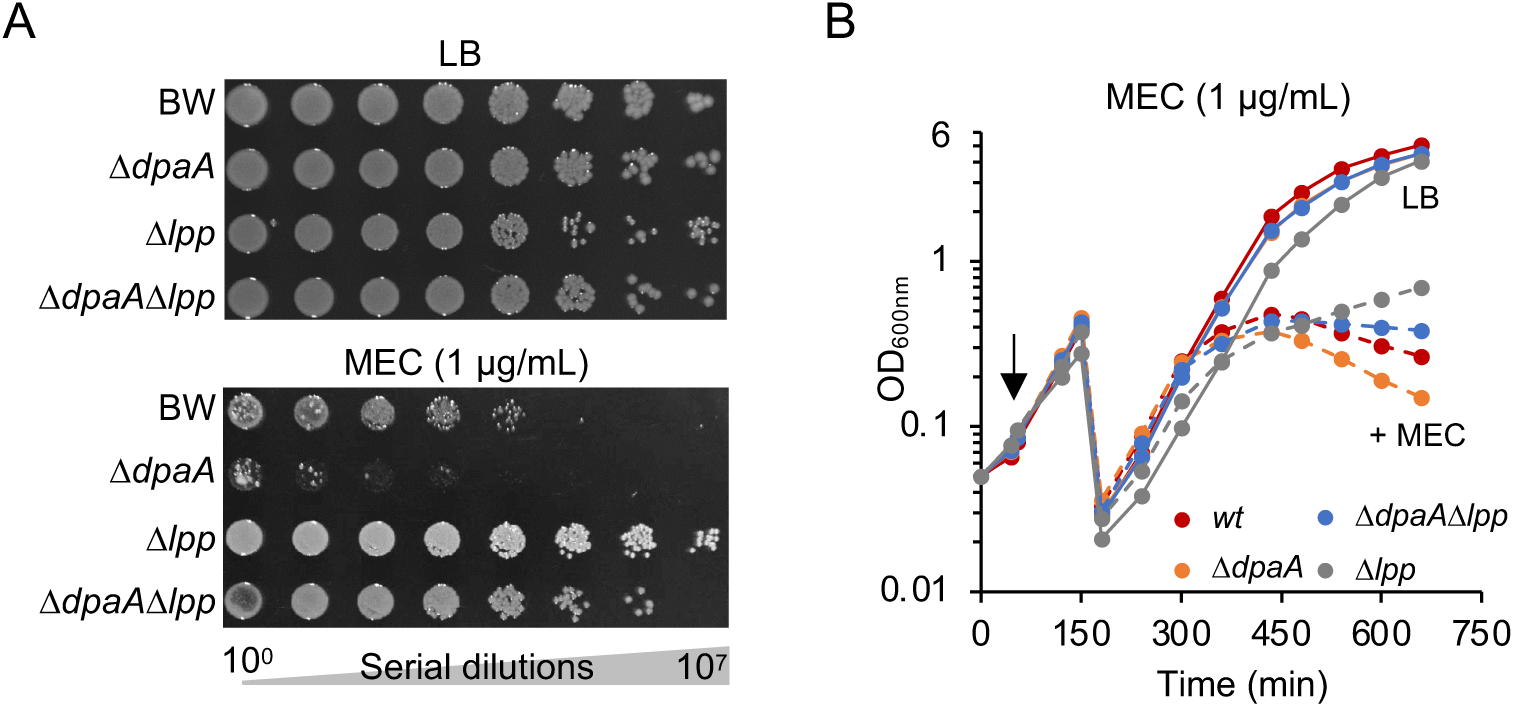
The *dpaA* mutant is more susceptible to mecillinam. (**A**) Serial dilutions of BW25113 and mutants (Δ*dpaA*, Δ*lpp* or Δ*dpaA*Δ*lpp*) were spotted on LB plates with or without mecillinam. Images were taken after incubation at 30°C for 33 h. (**B**) Growth of the same strains in liquid medium in the absence of mecillinam or upon addition of 1 µg/ml mecillinam (arrow) followed by measuring the optical density at 600 nm. Cultures were diluted 20-fold when the OD_600nm_ reached 0.5. Figure S6 shows that the effect is specific to mecillinam.

### DpaA is present in many bacteria that lack Lpp

We next asked if DpaA co-occurs with Lpp in diderm bacteria. We analysed the presence of proteins of the YkuD family (PF03734 in PFAM), Lpp-like proteins (PF04728) and DpaA-like proteins within the AnnoTree database, which contains a set of representative, fully-sequenced and consistently-annotated bacterial genomes (38). YkuD-like proteins are present in 72% of the genomes, often with several copies (61,365 sequences in 16,986 genomes). Lpp-like proteins are only present within a subset of γ-proteobacteria (∼3% of all genomes) (**Fig. 7A**). To identify DpaA-like proteins, we searched within the 61,365 YkuD-family sequences in AnnoTree using BLAST and the *E. coli* DpaA sequence as query, resulting in 2451 hits from 2310 different genomes (**Fig. 7A**). The identified sequences are present in a wide range of diderm bacteria lacking Lpp, including species within α-, γ- and δ-proteobacteria, Acidobacteria and Bacteriodetes (**Fig. 7A**). About a third of the genomes encoding Lpp did not contain a DpaA-like protein, suggesting that those organisms are not capable of detaching Lpp from PG or other proteins have this function (**Fig. 7A**). Alignment of the sequences of DpaA-like proteins identified within AnnoTree revealed the catalytic cysteine and several conserved positions common to all YkuD-like proteins (**Fig. 7B**). As in *E. coli* DpaA (**Fig. 1**), most DpaA-like proteins have alanine or asparagine 2 positions after the catalytic cysteine instead of an Arg that is present in LDTs with TPase activity.

**Figure 7.**
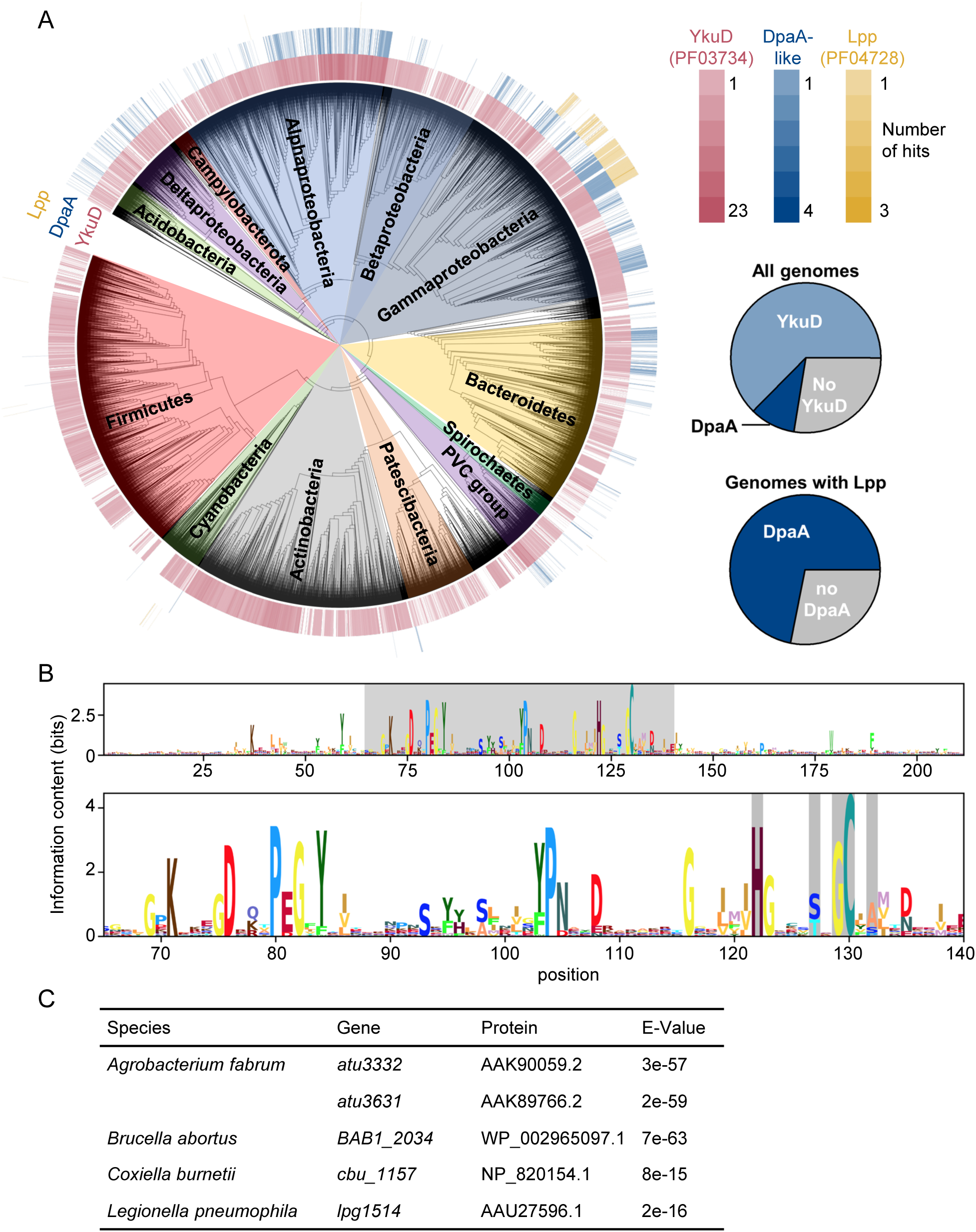
Conservation and distribution of DpaA. (**A**) Distribution of genes encoding proteins of the YkuD family (PFAM PF03734, red), Lpp (PF04728, yellow) and DpaA-like proteins (identified by BLAST within PF03734 proteins, blue) within the AnnoTree bacterial genome database. The intensity of the colour corresponds to the number of genes identified in each genome. The pie charts in the right bottom corner show the percentage of genomes in the database encoding proteins in the PF03734 family, DpaA-like proteins or none of these (left), and the percentage of PF04728-containing genomes also containing DpaA-like proteins. (**B**) Logogram obtained from the multiple sequence alignment of the DpaA-like proteins identified within the AnnoTree database. Top: full alignment; bottom: amino acid positions 65 to 140, highlighting same positions as in the alignment in Fig. 1A. (**C**) Table showing genes encoding DpaA-like proteins identified in bacteria lacking *lpp* and known to attach OM β-barrel proteins to PG. Proteins, NCBI protein annotation corresponding to the indicated gene; E-Value, BLAST *expect* value indicating the similarity to the query sequence, *E. coli* DpaA.

Overall, our sequence analysis suggests that species without Lpp use DpaA for a different purpose. Some of the DpaA homologues might function as LD-carboxypeptidase, consistent with their classification in the MEROPS peptidase database (39) within the C82.A01 peptidase subfamily along with the LD-carboxypeptidases Csd6 from *Helicobacter pylori* and Pgp2 from *Campylobacter jejuni* (40, 41). Alternatively, or in addition, DpaA could detach different substrates from PG, for example OMPs in α- and γ-proteobacteria (24, 25). Indeed, a search within the genomes of α- and γ-proteobacteria with PG-attached OMPs identified genes encoding DpaA-like proteins in all of them (**Fig. 7C**).

## DISCUSSION

DpaA is the first member of the YkuD family that does not act on LD-peptide bonds and is not a transpeptidase. Our work demonstrated that DpaA hydrolyses the amide bond between the L-centre of *m*DAP in PG and the ε-amino group of Lpp (**Fig. 8**). Hence, the name LdtF (LD-transpeptidase F) was not appropriate anymore and we propose to name the enzyme as Peptidoglycan *m*<u>D</u>ap-<u>p</u>rotein <u>a</u>midase <u>A</u> (DpaA).

**Figure 8.**
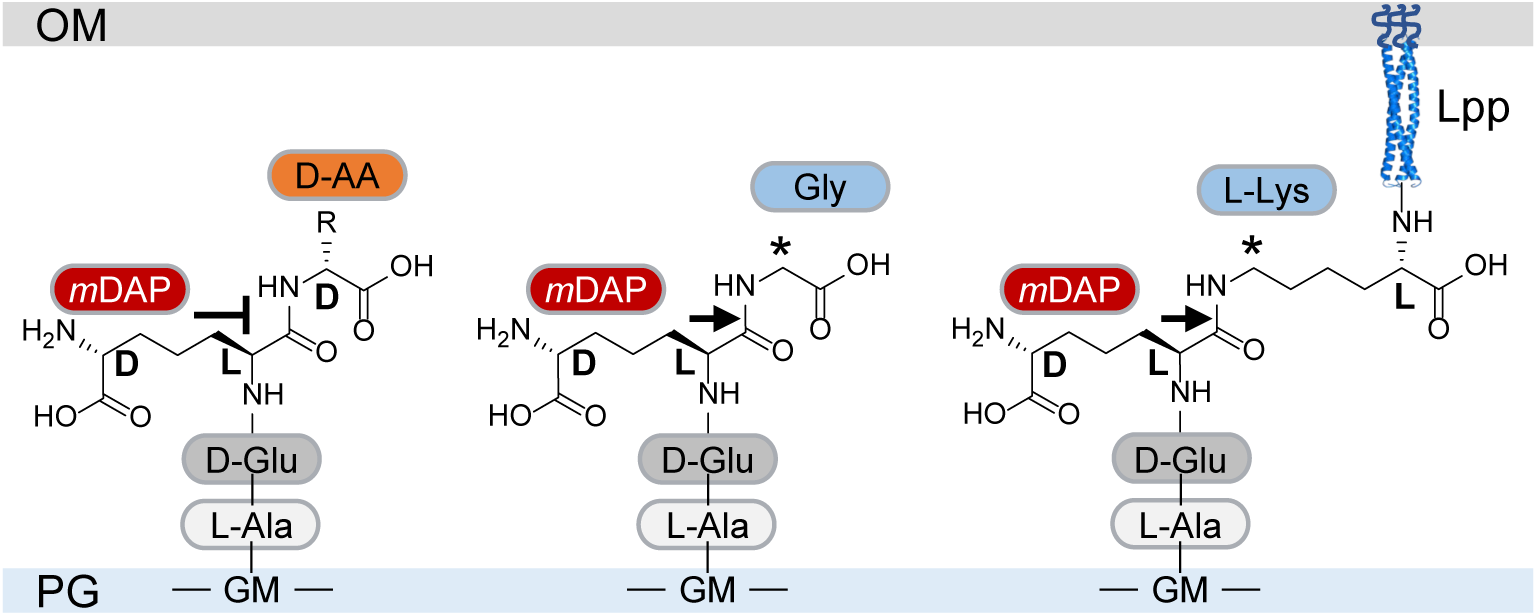
Specificity of DpaA. DpaA hydrolyses the amide bond between the L-centre of *m*DAP and Gly (middle) or the L-centre of mDAP and the ε-amino group of the C-terminal L-lysine of Lpp. Both substrates have a CH_2_ group adjacent to the cleavage site (labelled with *).

Lpp is the most abundant protein in *E. coli* and its attachment to PG substantially contributes to the stability of the cell envelope and prevents excessive OM vesiculation (5, 6, 9, 17). Why does the cell then have an enzyme that detaches Lpp from PG? We do not yet fully understand the importance of Lpp-detachment at different conditions. Our initial data indicate that DpaA might be required under certain stress conditions. The *dpaA* mutant has reduced fitness when cells grow in the presence of mecillinam, an antibiotic that induces a stark change in cell shape (from rod-shape to spherical shape) in growing cells (35). We hypothesize that under these conditions the detachment of Lpp from PG provides flexibility to the cell envelope to better coordinate the remodelling of the PG and OM during the shape change. The detachment of Lpp from PG could also be a means of controlling the amount of OM-derived vesicles during an infection, affecting the interaction of a diderm bacterium with the host organisms (42). In addition, the amount of PG-bound Lpp could affect bacterial cells within a biofilm and the outcome of bacterial competition (42b). However, it remains to be studied in more detail if DpaA affects virulence and bacterial fitness under environmental conditions or cell envelope stress.

DpaA becomes essential when the export of LPS to the outer membrane is compromised (29), presumably due to the spurious activation of peptidoglycan amidases via ActS (Gurnani *et al., under review*). Our TraDIS data revealed another connection between LPS and DpaA. The *lapAB* genes were interrupted by the transposon at much higher frequency in the *dpaA* mutant than the wild-type. Deleting *lapB* stabilizes LpxC and deregulates (enhances) LPS biosynthesis (43, 44), which causes lethality in wild-type cells but not in a hypervesiculating *lpp* mutant, which presumably releases the excess of LPS into outer membrane derived vesicles (30). The fact that transposon insertions in *lapAB* are recovered at a higher frequency in the *dpaA* mutant when compared to the parent strain is consistent with the lower fitness of the *dpaA* mutant in the presence of CHIR-090 (34), an inhibitor of LpxC, and suggests that enhanced PG-bound Lpp (due to the absence of DpaA) has a negative effect on LPS synthesis and/or export. We hypothesize that the higher amount of PG-bound Lpp in the *dpaA* mutant hinders (but not completely abolishes) LPS export, which alleviates problems caused by excessive LPS biosynthesis. This effect of the increased amount of PG-attached Lpp points to a yet unrecognized regulatory function of Lpp in LPS export.

Structure-based sequence alignment shows that LD-carboxypeptidases and DpaA-like proteins are closely related and that both lack the conserved arginine residue present in LD-TPases (**Fig. S7**). However, some residues conserved in DpaA-like proteins at the start of the catalytic domain are absent in the LD-carboxypeptidases (**Fig. S7**), and a BLAST search with *E. coli* DpaA failed to identify Csd6 while Pgp2 was identified with a low expect value (1e-4) (40, 41). Consistent with this low similarity, *E. coli* DpaA was not able to remove terminal D-alanine residues from PG stem peptides, however, more experiments will be necessary to determine whether some DpaA-like proteins show LD-carboxypeptidase activity.

Depending on the growth medium, *E. coli* naturally contains a small proportion of stem peptides with glycine residues at position 4 (14). Other bacteria use LDTs to incorporate non-canonical D-amino acids produced by themselves or present in the environment into the same position (45, 46). Of the 4 amino acids tested, DpaA was only capable of removing glycine residues from position 4, but not the three D-amino acids tested. However, at present we cannot exclude the possibility that some bacteria employ DpaA-like enzymes to remove non-canonical D-amino acids, which are toxic when incorporated at a high amount (47). The recent discovery of PG-attached OMPs in certain α-proteobacteria suggests that the DpaA-like enzymes in these species function to detach OMPs from PG (24, 25). In support of this hypothesis, the attachment of OMPs to PG occurs via N-terminal glycine or alanine, and the attachment of OM-lipoprotein LimB via an internal lysine residue, producing the very same amide bonds as in TetraGly4 or in PG-bound Lpp in *E. coli*. Moreover, DpaA homologues are present in many bacteria, including many species that do not contain Lpp, suggesting different substrates. Hence, it remains to be tested if DpaA-like enzymes are capable of removing OMPs from PG.

In summary, our work discovered that DpaA performs a yet unknown reaction in the bacterial cell envelope, the hydrolytic removal of a protein from PG. Presumably, DpaA provides flexibility to the bacterial cell to remodel the cell envelope when coping with certain stress situations. The discovery of this reaction also illustrates that cell envelope remodelling contributes to the robustness of bacterial cells and their lifestyles.

## MATERIALS AND METHODS

### Bacterial strains and growth conditions

Strains used in this study are listed in Table S2. Bacteria were grown aerobically at 30°C or 37°C on LB-plates or in liquid LB medium (10 g/L tryptone, 5 g/L yeast extract, 10 g/L NaCl; 15 g/L agar for plates). Antibiotics were used at the following concentrations: chloramphenicol (Cam), 25 µg/mL; kanamycin (Kan), 50 µg/mL; tetracycline (Tet), 5 µg/mL. For monitoring cell cultures treated with mecillinam, 30°C pre-warmed LB medium was inoculated with overnight cultures starting with a normalised OD_600nm_ of 0.05. The cultures were split in half at an OD_600nm_ of 0.1. One-half was supplemented with 1 µg/mL mecillinam, the other half remained untreated. The cultures were diluted 20-fold in fresh 30°C pre-warmed LB supplemented with and without mecillinam at an OD_600nm_ of 0.5.

### Construction of *E. coli* deletion strains

Deletion strains were obtained by transducing *kan*-marked alleles from the Keio *E. coli* single-gene knockout library (48) by P1 phage (49). The *kan* cassette was removed by pCP20-encoded Flp recombinase to generate deletions with a FRT-site scar sequence (50). The removal of the *kan* gene was verified by polymerase chain reaction. Strains with multiple deletions were generated by sequential P1 transduction and *kan* cassette removal.

### Plasmid construction

Plasmids and oligonucleotides used in this study are shown in Tables S2 and S3, respectively. Genes were amplified by PCR using DNA from MC1061 as template. pET28a-*ldtE* was constructed by ligase-independent cloning (51). Site directed mutagenesis of plasmids were performed using Q5^®^ Site-Directed Mutagenesis Kit (New England BioLabs) following manufacturer’s instructions.

For pET29b-DpaA-his construction the signal sequence of *dpaA* was substituted with the one of *pelB* by a three-step PCR method (52) using pET27b as template for *pelB*ss. The chimeric *pelB*ss-*dpaA* gene was cloned in pET29b between the *Nde*I and *Nco*I restriction sites. The thrombin cleavable sequence was inserted in the linker region between the 3’ end of the gene and the sequence codifying the His-tag using the DpaA_pET29b_R primer.

### Spot plate assay

The spot plate assays were performed as described (33). Serial dilutions of bacterial cultures were spotted on LB plates supplemented with the appropriate antibiotics (Tet, 5 µg/mL; Cam, 25 µg/mL). The plates were incubated at 30°C for 33 h and images were taken with an InGenius Syngene Bio Imaging system.

### Mecillinam susceptibility testing

Minimal inhibitory concentration (MIC) of mecillinam towards BW25113 and mutants with single or multiple gene deletions were determined with MIC stripes (mecillinam) from Liofilchem (Roseto degli Abruzzi, Italy). Growing cells from cultures with an OD_600nm_ of 0.5 were harvested by centrifugation (13,000 × g, 5 min), washed three times with 0.9% NaCl, normalised to an OD_600nm_ of 0.125 and distributed onto LB-plates with cotton swabs. Plates contained tetracyclin (5 µg/mL) when used for strains with pTB63. An MIC-mecillinam test stripe was applied and the plates were incubated at 30°C or 37°C. Images were taken 24 h post incubation with an InGenius Syngene Bio Imaging system.

### PG isolation and analysis

PG was isolated from *E. coli* cells and analysed by reversed-phase HPLC as described (14). PG for DpaA activity assays containing bound Lpp was isolated as described (14) with the omission of treatments with α-amylase and pronase E.

### Purification of LdtE

*E. coli* LOBSTR-BL21(DE3) cells (Kerafast) harbouring the pET28a-LdtE-his plasmid were grown in 2 L of LB-autoinduction medium (LB supplemented with 0.5% glycerol, 0.05% glucose, 0.2% Lactose, for 20 h at 30°C (53). Cells were harvested by centrifugation for 15 min at 5,000 × g at 4°C. The cell pellet was resuspended in 100 mL of buffer I (20 mM HEPES/NaOH pH 7.5, 1 M NaCl, 1 mM DTT, 10% glycerol) supplemented with 1 mM phenylmethyl sulfonyl fluoride (Sigma Aldrich), 1 x protease inhibitor cocktail (Sigma Aldrich) and desoxyribonuclease I (Sigma Aldrich). Cells were broken by sonication and insoluble cell debris was removed by ultracentrifugation for 1 h at 130,000 × g at 4°C. The supernatant was applied to a 5 mL HisTrap HP column equilibrated with buffer I on an ÄKTA PrimePlus. The column was washed with 20 mL of buffer I containing 40 mM imidazole and bound protein was eluted with buffer II (20 mM HEPES/NaOH pH 7.5, 500 mM NaCl, 1 mM DTT, 10% glycerol, 400 mM imidazole). Fractions containing LdtE were combined and dialysed against 3 L of buffer III (20 mM HEPES/NaOH pH 7.5, 300 mM NaCl, 10% glycerol, 0.1 mM TCEP). The sample was concentrated to 5 mL with a Vivaspin 6 filter. The protein was then further purified by size exclusion chromatography on a HiLoad 16/60 Superdex 200 (GE Healthcare) column using buffer III and a flowrate of 1 mL/min. LdtE-containing fractions were combined and stored at -80°C.

### Purification of DpaA and DpaA(C143A)

*E. coli* LOBSTR-BL21(DE3) (Kerafast) containing pET29b-DpaA-his (encoding PelB-DpaA(20-246)-His_6_) or pET29b-DpaA(C143A)-his (PelB-DpaA(20-246,C143A)-His_6_; cysteine-143 replaced by alanine) were grown at 25°C overnight in 4 L of TB-autoinduction medium (54). Cells were harvested by centrifugation for 15 min, 5,000 x g at 14°C. The cell pellet was resuspended in 150 mL of buffer I (20 mM HEPES/NaOH pH 7.5, 500 mM NaCl, 2 mM MgCl_2_, 10% glycerol) supplemented with 1 mM phenylmethyl sulfonyl fluoride (Sigma Aldrich), 1 x protease inhibitor cocktail (Sigma Aldrich) and desoxyribonuclease I (Sigma Aldrich). Cells were broken by sonication and the insoluble fraction was removed by ultracentrifugation for 1 h at 130,000 × g at 4°C. The supernatant was applied to a 5 mL HisTrap HP column pre-equilibrated with buffer I using an ÄKTA PrimePlus. The column was washed with 20 mL buffer I with 40 mM imidazole and protein was eluted in a 12-mL gradient of from buffer I with 40 mM imidazole to buffer I with 400 mM imidazole. Fractions containing DpaA were combined and split into two pools. Pool 1 was dialysed against 3 L of buffer III (20 mM HEPES/NaOH pH 7.5, 300 mM NaCl, 10% glycerol, 10 mM EDTA), pool 2 was incubated with 10 units of thrombin (restriction grade, Novagen) to remove the His_6_-tag, during dialysis against 3 L buffer III. Proteins were further purified by size exclusion chromatography on a HiLoad 16/60 Superdex 200 (GE Healthcare) column using buffer III and a flowrate of 1 mL/min. The protein-containing fractions were aliquoted and stored at -80°C.

### Purification of LdtD

LdtD was expressed in *E. coli* LOBSTR-BL21(DE3) cells harbouring the overexpression plasmid pETMM82 (55) and purified as previously described (32).

### HPLC activity assay

DpaA activity assays were carried out in a total volume of 50 µL in 20 mM HEPES/NaOH pH 7.5 or 20 mM Na-acetate (pH 5.0), with 100 mM NaCl, 0.05% Triton X-100 (reduced), 0.1 mM TCEP, 10 µL substrate (muropeptides or PG), and 2 µM of protein. A control sample contained no protein. Purified non-reduced and desalted muropeptides (Tri-LysArg and TetraTri-LysArg) were a gift from J.-V. Höltje, MPI Tübingen, Germany). The reaction mixture was incubated overnight in a thermoshaker at 37°C and 900 rpm. The reaction was stopped by boiling the samples for 10 min at 100°C. Samples were reduced with sodium borohydride and muropeptides were separated by reversed-phase HPLC (14), except that the gradient was 90 min instead of 135 min for samples with purified muropeptides. For activity assays on PG, the reaction products were treated overnight with the muramidase cellosyl (0.5 µg/mL) at 37°C and 900 rpm in 80 mM sodium phosphate pH 4.8, and the released muropeptides were reduced and separated by HPLC as described above. Tri-LysArg, TetraTri-LysArg and the products generated by DpaA from these muropeptides were collected during the HPLC run and analysed by MS/MS as described (56).

### Amino acid exchange followed by DpaA reaction

The amino acid exchange reaction with glycine, D-valine, D-glutamine or D-lysine was performed in a final volume of 50 µL in 20 mM HEPES/NaOH pH 7.5; 100 mM NaCl; 0.1 mM TCEP; 10 mM MgCl_2_; 0.05% Triton X-100. Ten µL of muropeptides obtained from PG of BW25113Δ6LDT (∼50 µg) were incubated with glycine, D-valine, D-glutamine or D-lysine (10 mM) and LdtD (5 µM) for 6 h at 37°C. The reaction was terminated by boiling the sample for 10 min. The sample was centrifuged and the supernatant containing the soluble muropeptides were split into two equal aliquots. In one half, the pH was adjusted to 5.0 with Na-acetate, DpaA (5 µM) was added and the sample was incubated overnight at 37°C. The other half was treated the same way without the addition of DpaA. Samples were boiled for 10 min at 100°C, treated with sodium borohydride and muropeptides were separated by HPLC as described above.

### DpaA activity assay against PG-Lpp

DpaA or DpaA(C143A) (2 µM) was mixed with 10 µL of PG with attached Lpp from BW25113*ΔdpaA* in 20 mM HEPES/NaOH pH 7.5, 100 mM NaCl, 0.05% Triton X-100 (reduced) and 0.1 mM TCEP. Samples were incubated overnight in a thermoshaker with 900 rpm at 37°C. Soluble proteins were separated from PG by centrifugation (13,000 × g, 10 min, 4°C). The supernatant was collected, proteins were separated by SDS-PAGE and stained with Coomassie Brilliant Blue, or transferred to nitrocellulose for Western blotting and detection with an anti-Lpp antibody (kind gift from Jean-François Collet, UC Louvain, Brussels, Belgium).

### Analysis of DpaA conservation

We used the AnnoTree database to study the conservation of DpaA-like proteins within bacteria (38). This database contains a set of archaeal and bacterial genomes which are completely sequenced and consistently annotated. The sequences of proteins identified as belonging to PFAM families PF03734 (LD-transpepetidase catalytic domain, 61,365 sequences in 16,986 genomes) and PF04728 (Lipoprotein leucine-zipper or Lpp, 717 sequences in 694 genomes) were downloaded from the AnnoTree website. We searched with the PF03734 sequences for DpaA-like proteins using BLAST (57) and *E. coli* DpaA as query with an e-value cut-off of 1e-8. This search resulted in 2451 hits from 2310 genomes. The number of proteins of each type per genome were counted using custom python scripts and the generated datasets were represented along a phylogenetic tree of all genomes in AnnoTree using iTOL (58). The sequences for all identified DpaA-like proteins were aligned using Clustal Omega (59). The resulting alignment was uploaded to the Skyling website to generate a logogram by converting to a hidden Markov model after removing mostly empty columns in the alignment (60). Next, the matrix with position frequencies was downloaded and the final logogram was generated using the logomaker python package (61). The search for DpaA-like genes in organisms attaching β-barrel outer membrane proteins to PG was conducted using BLAST and the genomes analysed were: *Agrobacterium fabrum* str. C58 (ASM9202v1), *Brucella abortus* 2308 (ASM74219v1), *Coxiella burnetii* RSA 493 (ASM776v2), and *Legionella pneumophila* subsp. pneumophila str. Philadelphia 1 (ASM848v1).

### TraDIS method

Transposon mutant libraries were constructed by electrotransformation of the mini-Tn*5* transposon carrying a kanamycin resistance cassette (Epibio). Two technical replicates of each library were prepared for sequencing as described previously (62). Samples were sequenced using an Illumina MiSeq with 150 cycle v3 cartridges. Raw data are available at the European Nucleotide Archive (ENA) accession XXXX. Data were demultiplexed using the Fastx barcode splitter, to remove the barcode unique to each sample (63). The transposon was matched and trimmed allowing for 4 bp mismatch, and surviving reads were mapped to the BW25113 reference genome using bwa mem (accession CP009273.1). 2,877,112 and 4,664,621 reads were mapped for BW25113 and BW25113Δ*dpaA*, respectively. Data are available for viewing at the TraDIS-vault browser: https://tradis-vault.qfab.org/. The BioTraDIS analysis package (version 1.4.5) was used to calculate the log-fold-change in read depth between each gene in the control and *dpaA* transposon libraries, using a threshold of >2-fold change and a Q-value < 0.01 (64).

## ACKNOWLEDGEMENTS

We thank Jean-François Collet (Université catholique de Louvain) for the gift of anti-Lpp antibodies.

---

**Figure S1.**
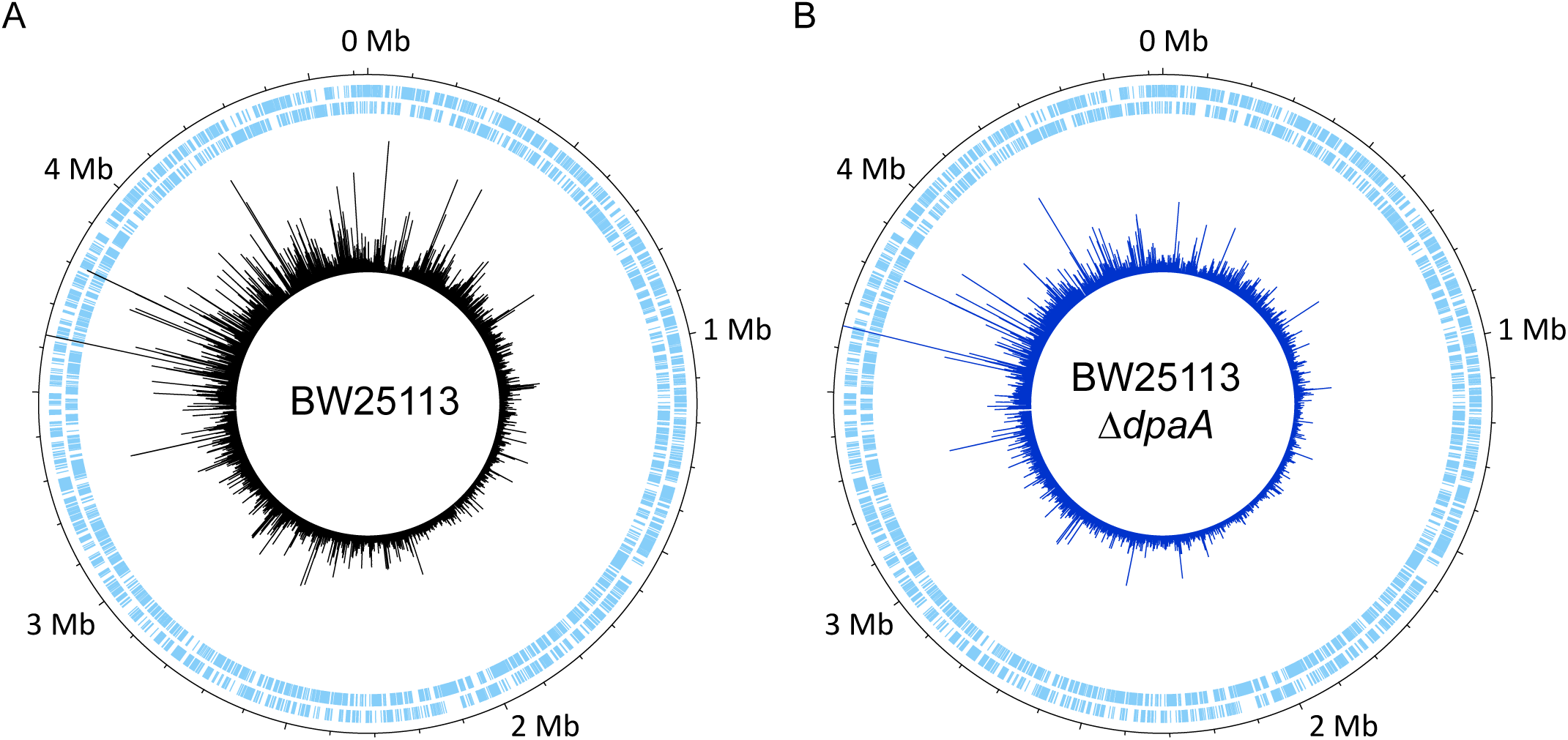
Construction of transposon libraries. Genome maps representing the transposon mutant libraries constructed in BW25113 (**A**) and BW25113Δ*dpaA* (**B**). The sense and anti-sense coding sequences are displayed in the two outermost tracks, respectively, in pale blue. The position and frequency of transposon insertion events around the BW25113 reference genome are represented by the peaks in the innermost track; plotted using DNAPlotter with a window size of 1 and a step size of 1.

**Figure S2.**
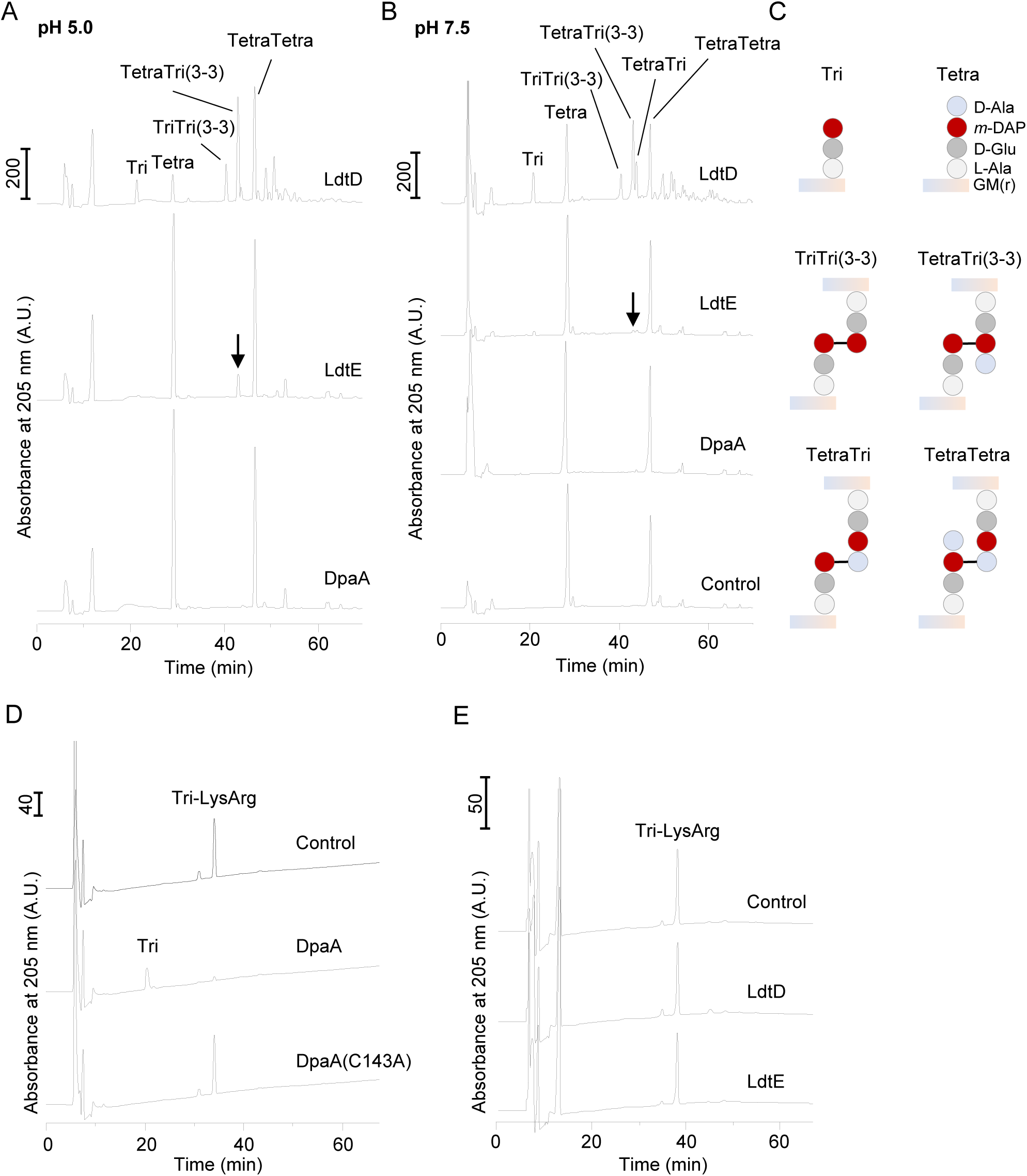
Distinct activities of LdtD/LdtE and DpaA. PG from BW25113Δ6LDT was incubated with LdtD, LdtE or DpaA at pH 5.0 (**A**) or pH 7.5 (**B**). The PG was then digested with cellosyl and the resulting muropeptides were separated by HPLC. The LD-TPase product TetraTri(3-3) was present in reactions with LdtD or LdtE, but not DpaA or the control without enzyme. (**C**) Structures of the relevant muropeptides. (**D**) Tri-LysArg incubated with DpaA, DpaA(C143A) and (**E**) LdtD or LdtE. Muropeptides were reduced with sodium borohydride and separated by HPLC. G, *N*-acetylglucosamine; M(r), *N*-acetylmuramitol; L-Ala, L-alanine; D-Glu, D-glutamic acid; D-Ala, D-alanine; *m*-DAP, *meso*-diaminopimelic acid.

**Figure S3.**
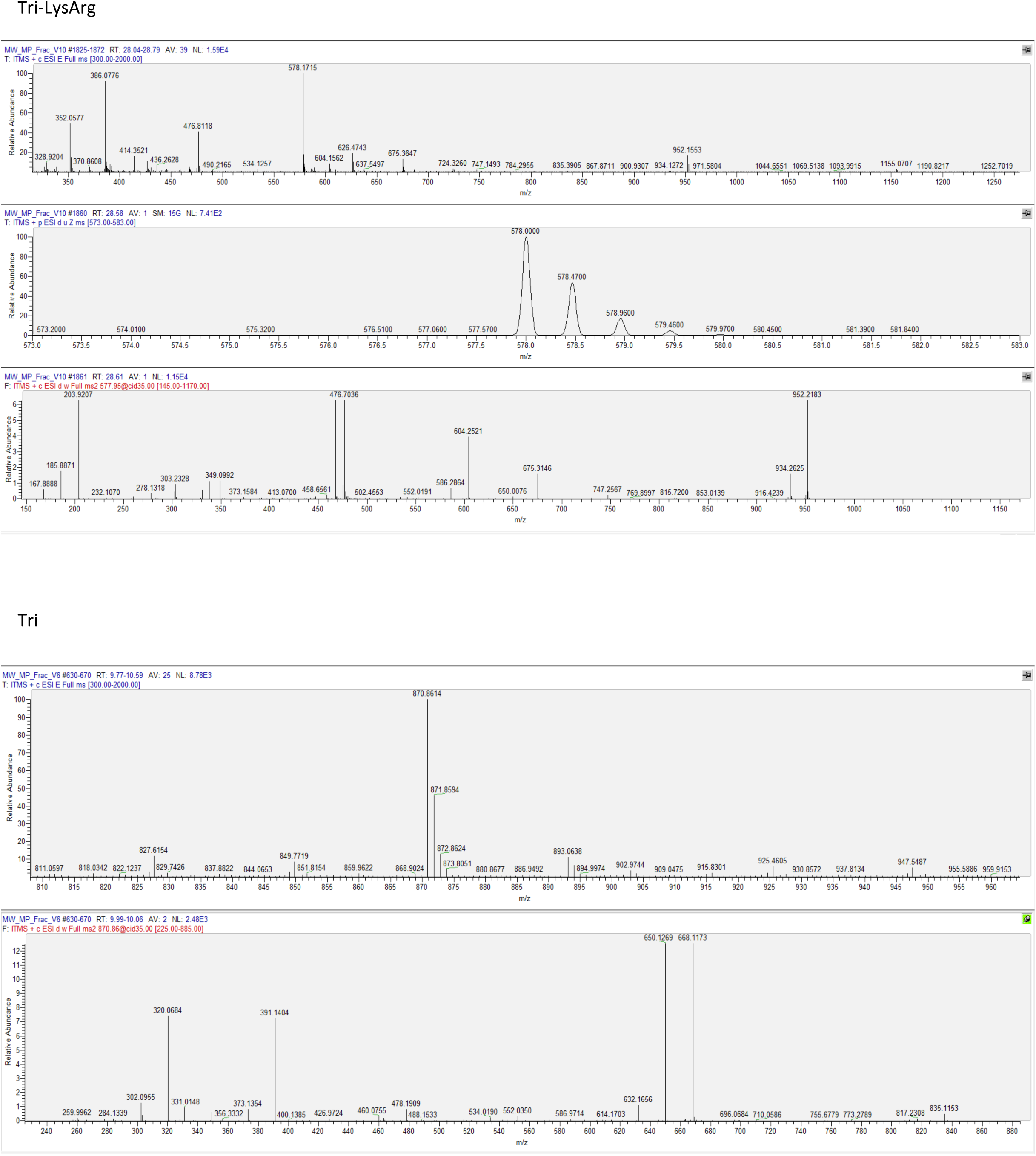
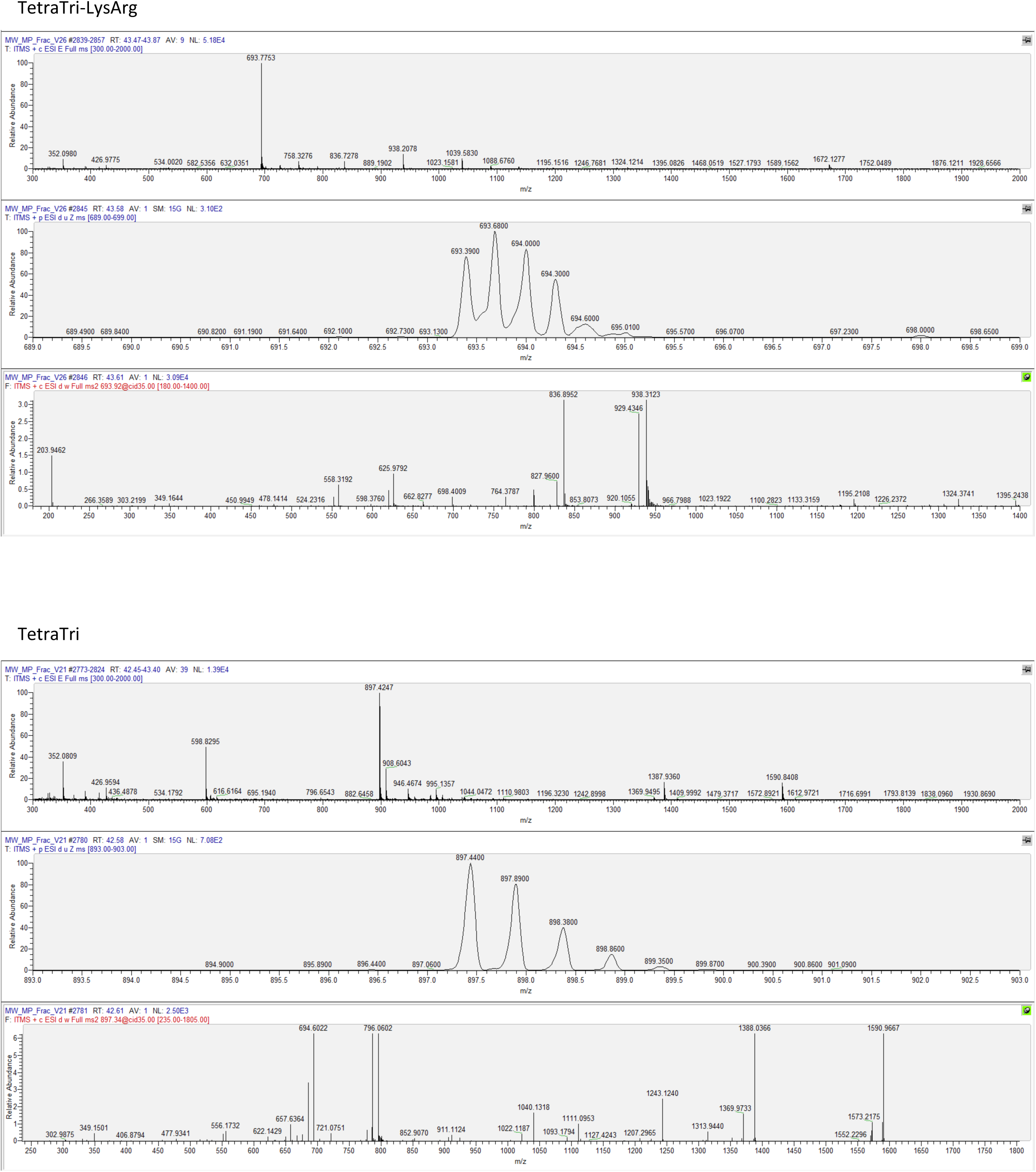
Mass spectrometry spectra of the substrates and products in DpaA reactions. Muropeptides (Tri-LysArg, Tri, TetraTri-LysArg, TetraTri) were manually collected during the HPLC runs (Fig. 3) and analysed by mass spectrometry (Bui *et al*., 2009). Spectra on the top show the unfragmented ions, the bottom MS/MS spectra were obtained after fragmentation. Middle spectra show zoomed regions of the relevant [M+nH]^n+^ ions to determine the charges. MS spectra confirmed the identities of the muropeptides analysed. <u>Reference:</u> Bui NK, Gray J, Schwarz H, Schumann P, Blanot D, Vollmer W. 2009. The peptidoglycan sacculus of *Myxococcus xanthus* has unusual structural features and is degraded during glycerol-induced myxospore development. *J Bacteriol* 191:494-505.

**Figure S4.**
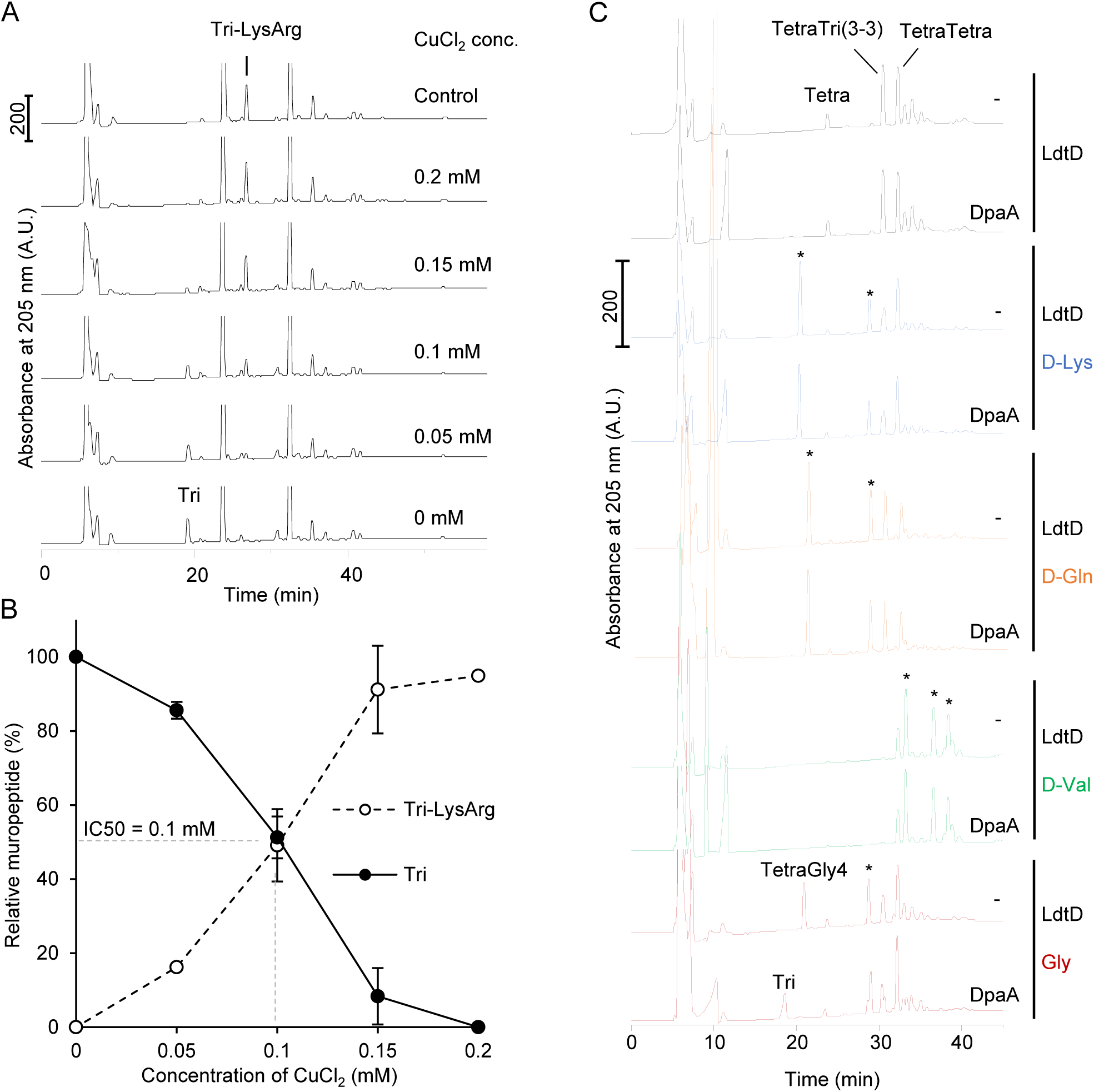
DpaA is inhibited by copper and does not release D-Lys, D-Val or D-Gln from PG. (**A**) PG from BW25113*ΔdpaA* was incubated with DpaA in the presence of increasing concentration of CuCl_2_. Muropeptides were generated, reduced and separated by HPLC. DpaA activity is seen as the reduction in the Tri-LysArg peak which decreased by copper in a concentration-dependent manner. Chromatographs were cropped above 250 mAU to better observe Tri-LysArg. (**B**) Quantification of the relative amount of Tri-LysArg and Tri plotted against the concentration of CuCl_2_. The data are mean ± variation of two independent experiments. (**C**) PG from BW25113Δ6LDT was incubated with LdtD and different D-amino acids, which were incorporated into position 4 of the stem peptides. A star (*) labels muropeptides with non-canonical D-amino acids. The PG was incubated with DpaA, followed by generation of muropeptides, reduction and HPLC analysis. DpaA released glycine from PG, but not the D-amino acids tested.

**Figure S5.**
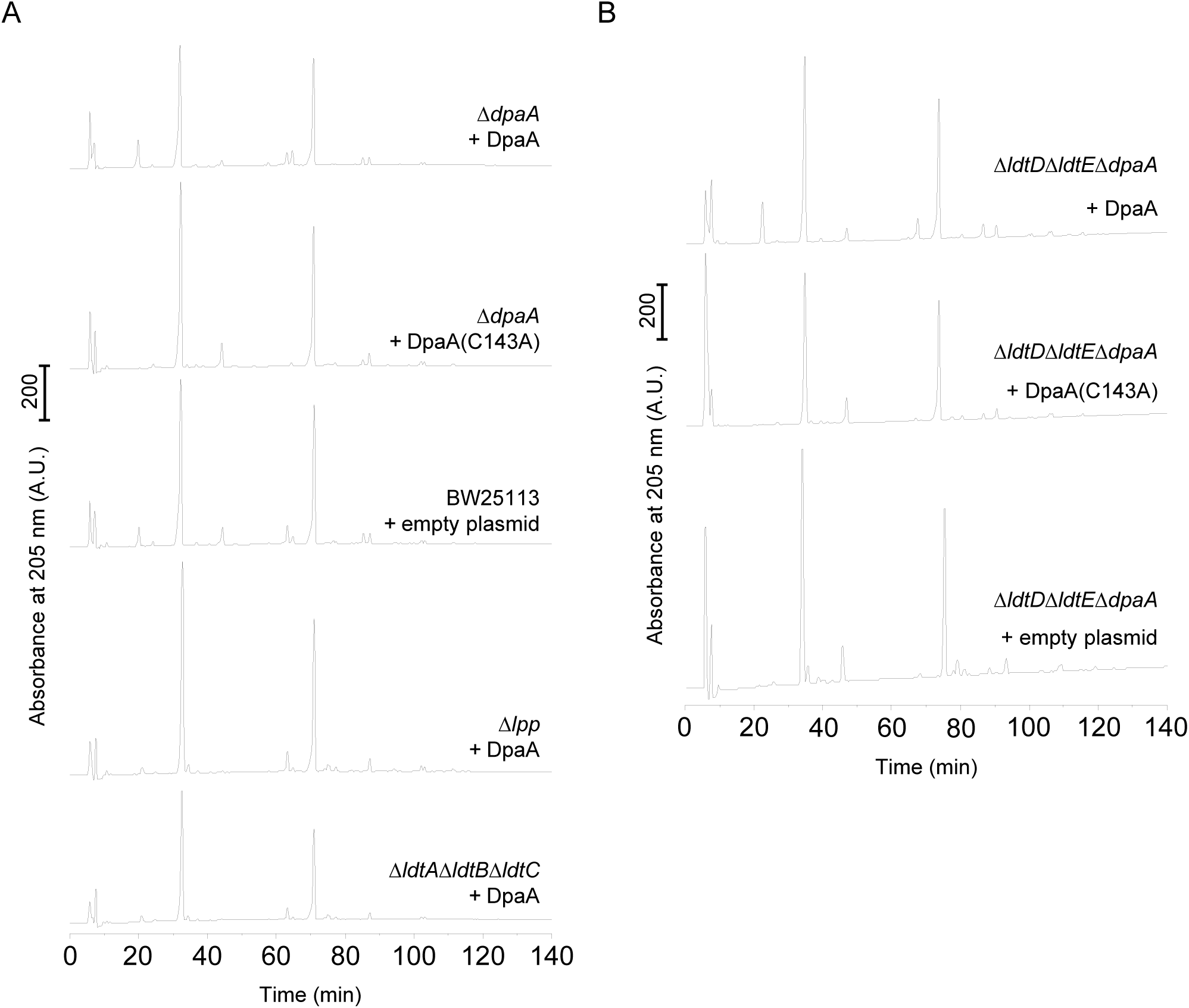
DpaA is active in cells. Full HPLC chromatograms of the cropped versions shown in Figure 5.

**Figure S6.**
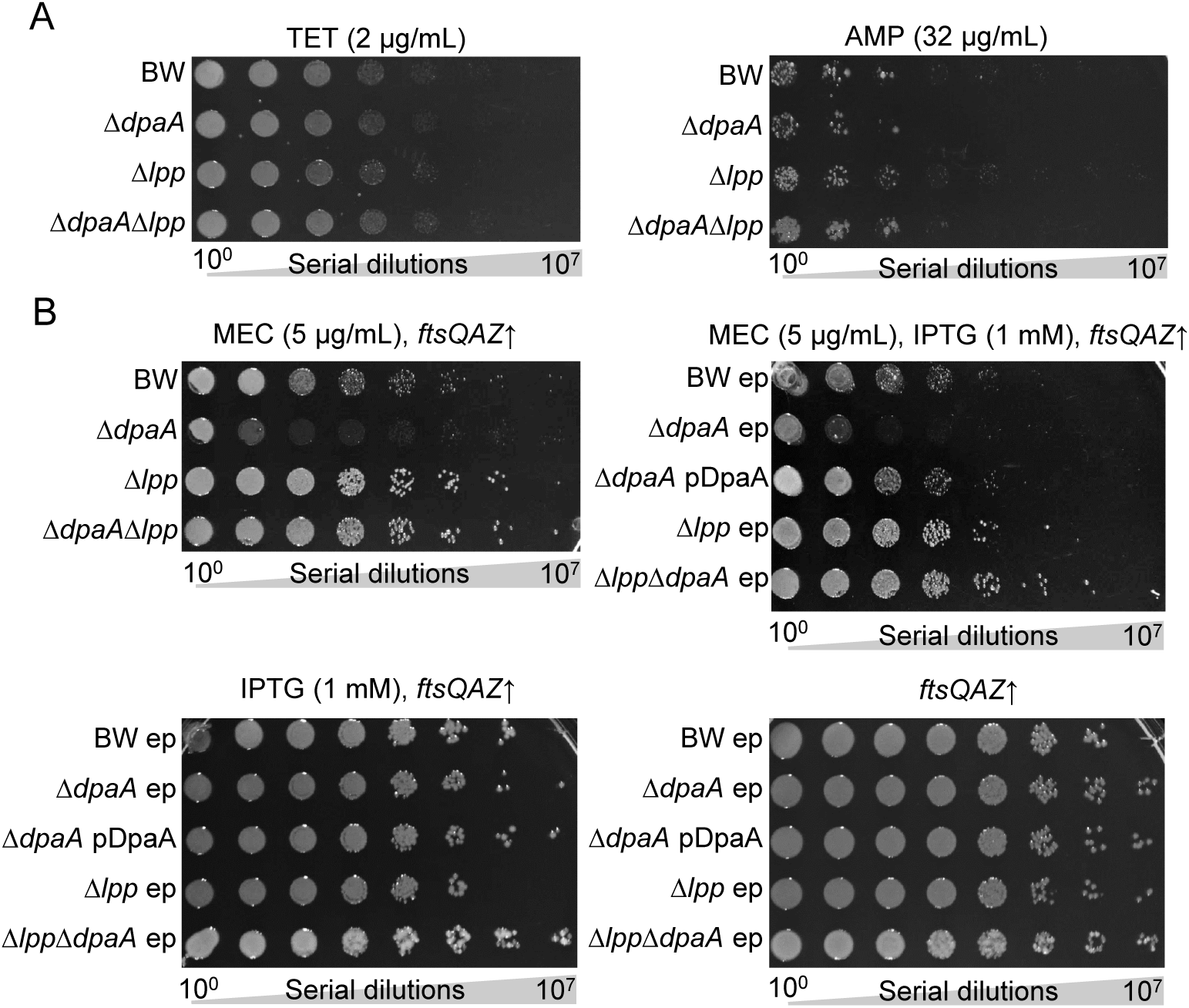
Growth defect of the *dpaA* mutant is specific to mecillinam. (**A**) Spot-plate assays with tetracycline (TET) or ampicillin (AMP) showed no altered sensitivity of the *dpaA* mutant. (**B**) Spot-plate assay of strains carrying the pTB63 plasmid, expressing *ftsQAZ* alone or in combination with empty plasmid (pGS100, ep) or plasmids expressing DpaA, in the presence or absence of mecillinam. Cells overexpressing *ftsQAZ* were more susceptible to mecillinan when they lacked *dpaA*.

**Figure S7.**
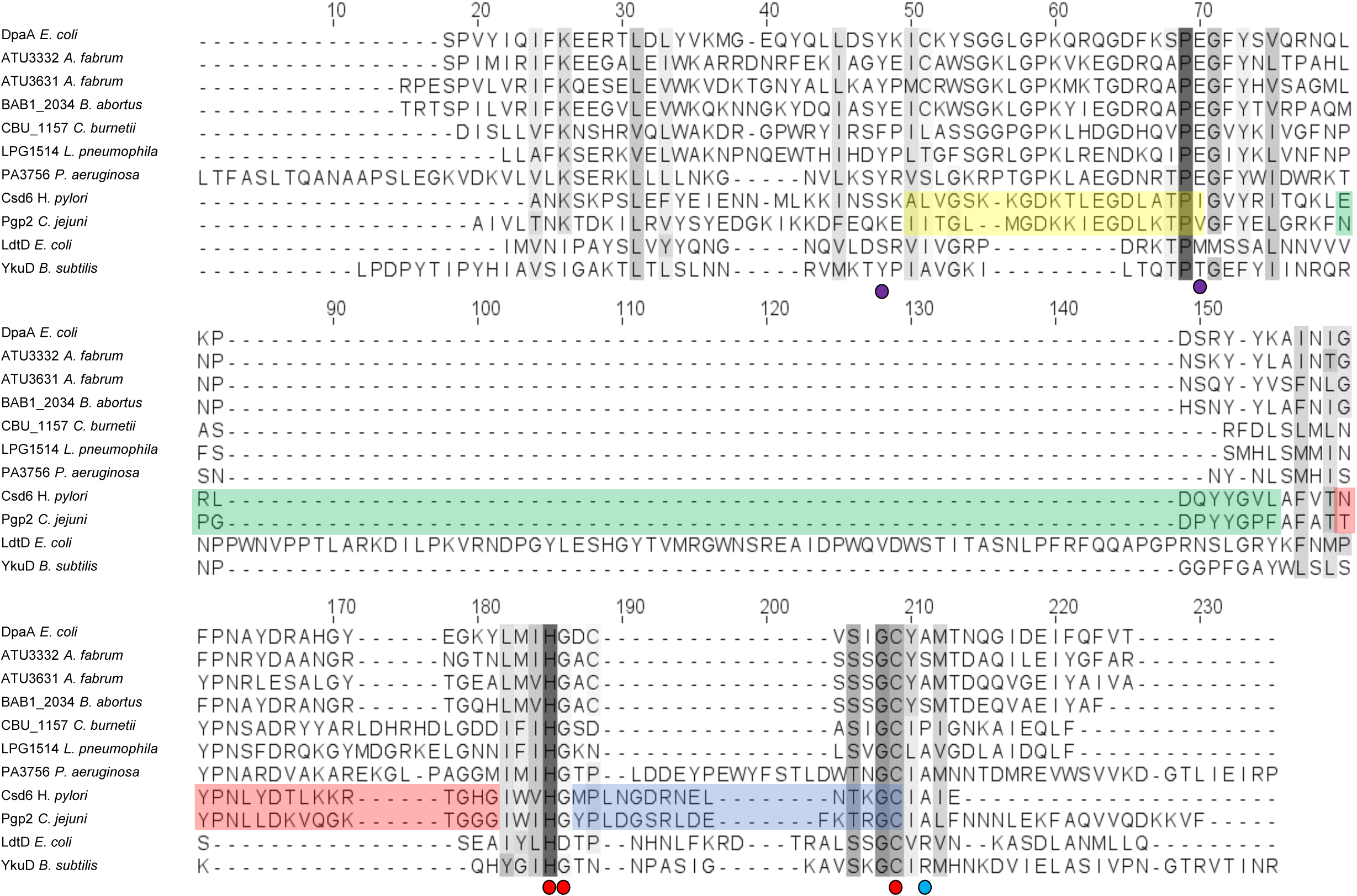
Sequence alignment of DpaA-like proteins and members of the YkuD family. Structure-based sequence alignment of DpaA-like proteins, LD-carboxypeptidases and LD-transpeptidases obtained with PROMALS3D (Pei *et al*., 2008), using the structures of *E. coli* LdtD (PDB accession: 6NTW), Bacillus subtilis YkuD (1Y7M) and *Helicobacter pylori* Csd6 (4XZZ) as guides. The residues corresponding to the 4 structural loops surrounding the substrate-pocket in Csd6 are highlighted in different colors. The Red dots indicate the catalytic triad in Csd6, purple dots mark positions conserved in DpaA-like proteins but different in LD-carboxypeptidases and the blue dot indicates the conserved Arg in LD-TPases but not in hydrolytic YkuD-family proteins. The sequences aligned include *E. coli* DpaA (Uniprot P0AA99, residues 45-159), *Agrobacterium fabrum* ATU3332 (Q7CS68, 43-168) and ATU3631 (A0A3G2CVN4, 58-176), *Brucella abortus* BAB1_2034 (C4ITM3, 50-169), *Coxiella burnetii* CBU_1157 (Q83CG1, 67-183), *Legionella pneumophila* LPG1514 (Q83XL0, 72-186), *Pseudomonas aeruginosa* PA3756 (Q9HXN9, 10-166), *Helicobacter pylori* Csd6 (O25255, 73-181) and *Campylobacter jejuni* Pgp2 (A7H3E2, 66-196), *E. coli* LdtD (P22525,377-543) and *Bacillus subtilis* YkuD (O34816, 48-164). <u>Reference:</u> Pei J, Kim B-H, Grishin NV. 2008. PROMALS3D: a tool for multiple protein sequence and structure alignments. *Nucleic Acids Research* 36:2295-2300.

**Table S1.** TraDIS raw data.

**Table S2.**
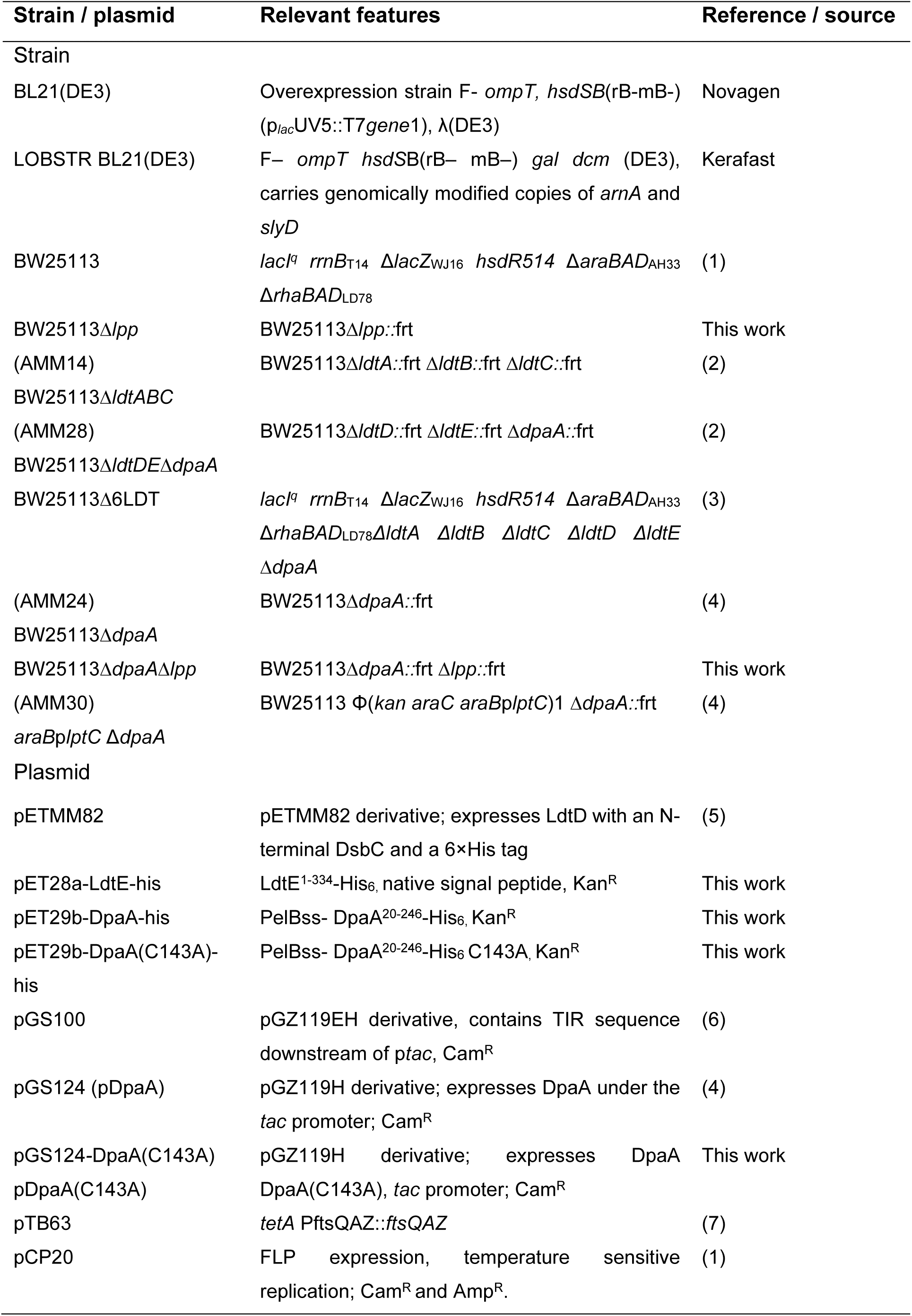
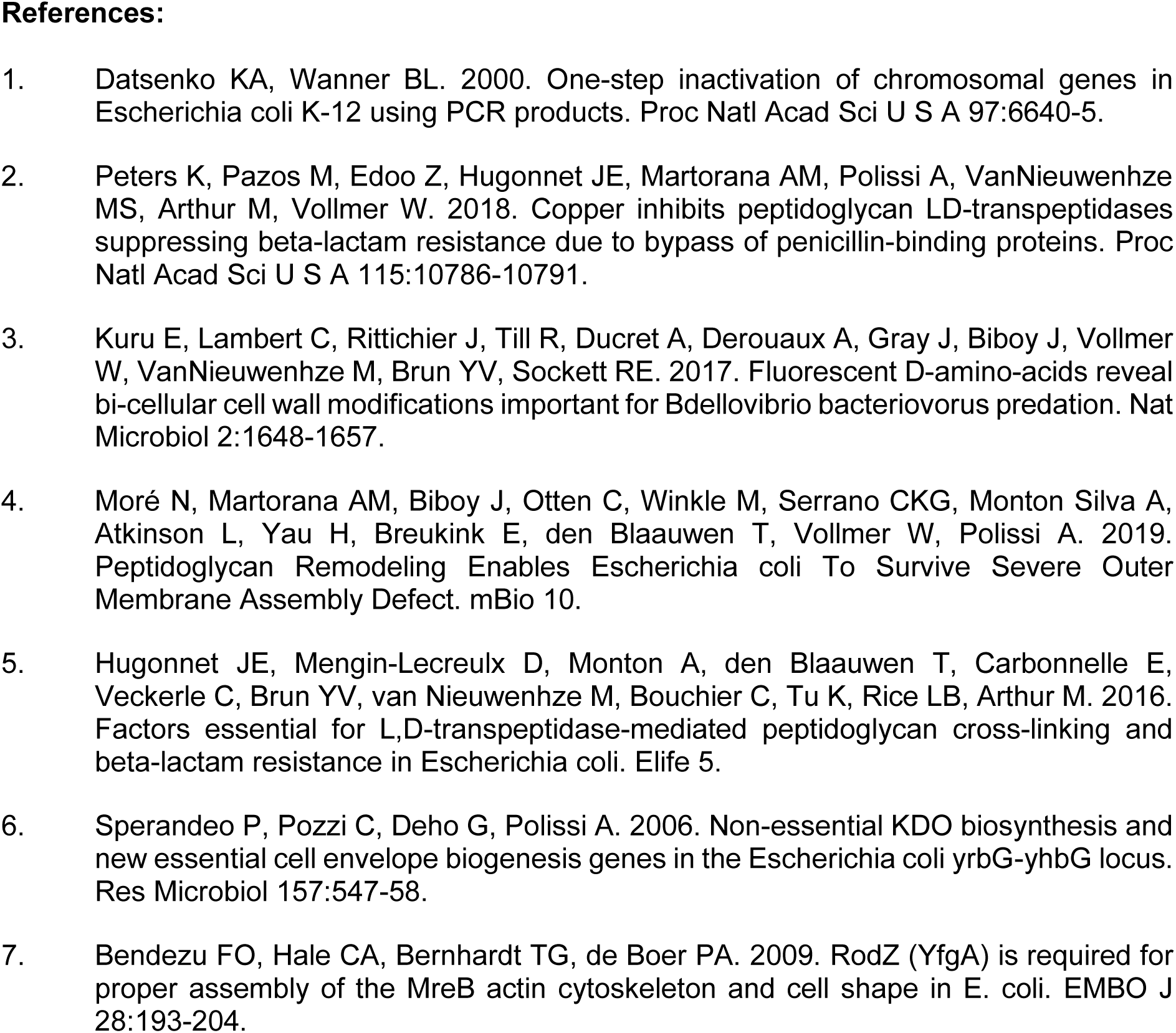
Strains and plasmids used in this work.

**Table S3.**
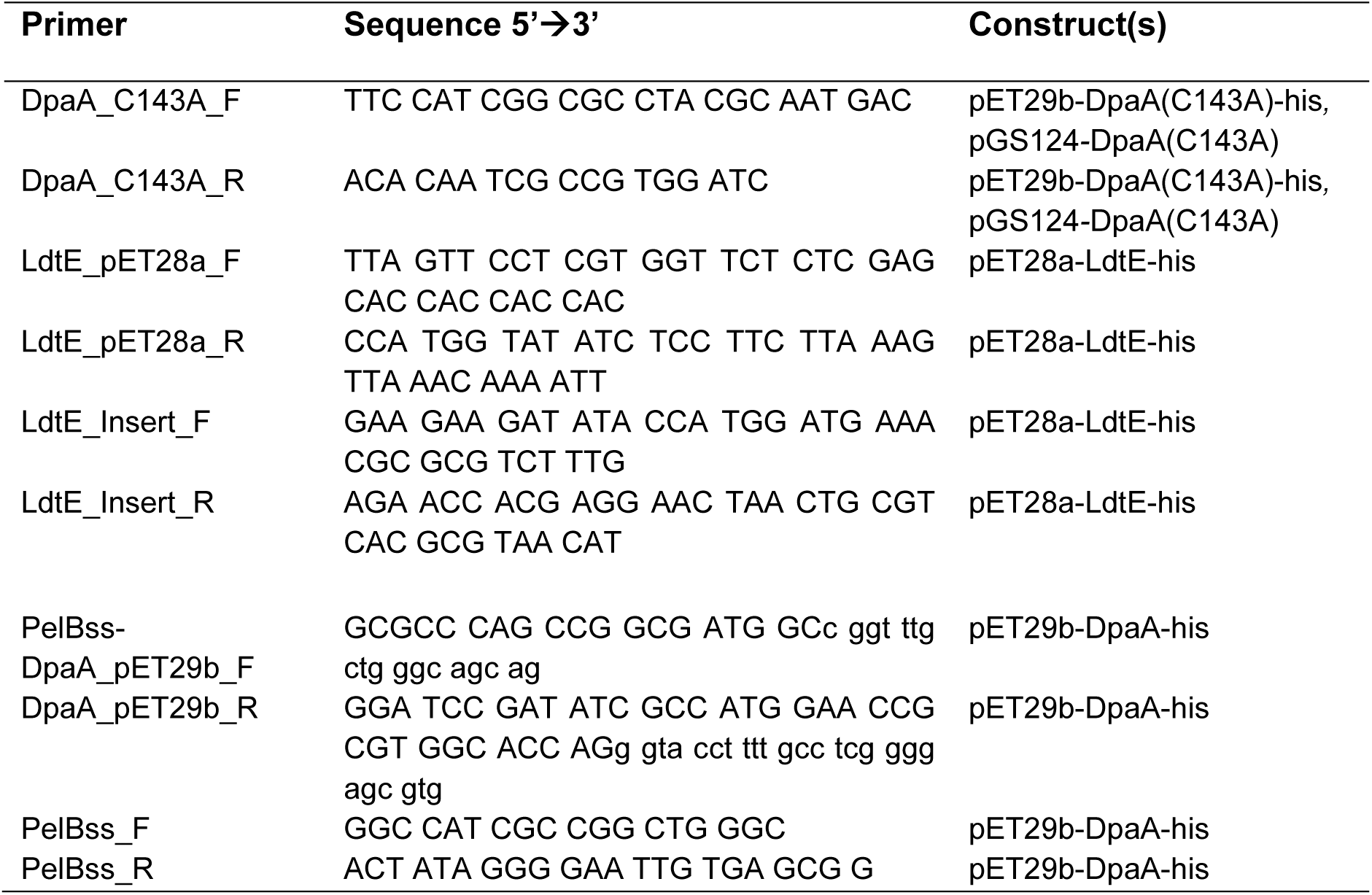
Oligonucleotides used in this study.

## REFERENCES

1. Silhavy TJ, Kahne D, Walker S. 2010. The bacterial cell envelope. Cold Spring Harb Perspect Biol 2:a000414. doi: 10.1101/cshperspect.a000414

2. Vollmer W, Blanot D, de Pedro MA. 2008. Peptidoglycan structure and architecture. FEMS Microbiol Rev 32:149–67. doi.org/10.1111/j.1574-6976.2007.00094.x

3. Egan AJF, Errington J, Vollmer W. 2020. Regulation of peptidoglycan synthesis and remodelling. Nat Rev Microbiol 18:446–460. doi.org/10.1038/s41579-020-0366-3

4. Braun V, Rehn K. 1969. Chemical characterization, spatial distribution and function of a lipoprotein (murein-lipoprotein) of the *E. coli* cell wall. The specific effect of trypsin on the membrane structure. Eur J Biochem 10:426–38. doi: 10.1111/j.1432-1033.1969.tb00707.x

5. Li GW, Burkhardt D, Gross C, Weissman JS. 2014. Quantifying absolute protein synthesis rates reveals principles underlying allocation of cellular resources. Cell 157:624–35. doi: 10.1016/j.cell.2014.02.033

6. Cascales E, Bernadac A, Gavioli M, Lazzaroni JC, Lloubes R. 2002. Pal lipoprotein of *Escherichia coli* plays a major role in outer membrane integrity. J Bacteriol 184:754–9. doi: 10.1128/JB.184.3.754-759.2002

7. Parsons LM, Lin F, Orban J. 2006. Peptidoglycan recognition by Pal, an outer membrane lipoprotein. Biochemistry 45:2122–8. doi.org/10.1021/bi052227i

8. Suzuki H, Nishimura Y, Yasuda S, Nishimura A, Yamada M, Hirota Y. 1978. Murein-lipoprotein of *Escherichia coli*: a protein involved in the stabilization of bacterial cell envelope. Mol Gen Genet 167:1–9. doi.org/10.1007/BF00270315

9. Yem DW, Wu HC. 1978. Physiological characterization of an *Escherichia coli* mutant altered in the structure of murein lipoprotein. J Bacteriol 133:1419–26. doi: 10.1128/JB.133.3.1419-1426.1978

10. Hantke K, Braun V. 1973. Covalent binding of lipid to protein. Diglyceride and amide-linked fatty acid at the N-terminal end of the murein-lipoprotein of the *Escherichia coli* outer membrane. Eur J Biochem 34:284–96. doi: 10.1111/j.1432-1033.1973.tb02757.x

11. Okuda S, Tokuda H. 2011. Lipoprotein sorting in bacteria. Annu Rev Microbiol 65:239–59. doi.org/10.1146/annurev-micro-090110-102859

12. Braun V. 1975. Covalent lipoprotein from the outer membrane of *Escherichia coli*. Biochim Biophys Acta 415:335–77. doi: 10.1016/0304-4157(75)90013-1.

13. Magnet S, Bellais S, Dubost L, Fourgeaud M, Mainardi JL, Petit-Frere S, Marie A, Mengin-Lecreulx D, Arthur M, Gutmann L. 2007. Identification of the L,D-transpeptidases responsible for attachment of the Braun lipoprotein to *Escherichia coli* peptidoglycan. J Bacteriol 189:3927–31. doi: 10.1128/JB.00084-07

14. Glauner B, Holtje JV, Schwarz U. 1988. The composition of the murein of *Escherichia coli*. J Biol Chem 263:10088–95. doi.org/10.1016/S0021-9258(19)81481-3

15. Asmar AT, Ferreira JL, Cohen EJ, Cho SH, Beeby M, Hughes KT, Collet JF. 2017. Communication across the bacterial cell envelope depends on the size of the periplasm. PLoS Biol 15:e2004303. doi.org/10.1371/journal.pbio.2004303

16. Cohen EJ, Ferreira JL, Ladinsky MS, Beeby M, Hughes KT. 2017. Nanoscale-length control of the flagellar driveshaft requires hitting the tethered outer membrane. Science 356:197–200. doi: 10.1126/science.aam6512

17. Hoekstra D, van der Laan JW, de Leij L, Witholt B. 1976. Release of outer membrane fragments from normally growing *Escherichia coli*. Biochim Biophys Acta 455:889–99. doi: 10.1016/0005-2736(76)90058-4

18. Hirota Y, Suzuki H, Nishimura Y, Yasuda S. 1977. On the process of cellular division in *Escherichia coli*: a mutant of *E. coli* lacking a murein-lipoprotein. Proc Natl Acad Sci U S A 74:1417–20. doi: 10.1073/pnas.74.4.1417

19. Uhlich GA, Gunther NWt, Bayles DO, Mosier DA. 2009. The CsgA and Lpp proteins of an *Escherichia coli* O157:H7 strain affect HEp-2 cell invasion, motility, and biofilm formation. Infect Immun 77:1543–52. doi: 10.1128/IAI.00949-08

20. Sha J, Fadl AA, Klimpel GR, Niesel DW, Popov VL, Chopra AK. 2004. The two murein lipoproteins of *Salmonella enterica serovar Typhimurium* contribute to the virulence of the organism. Infect Immun 72:3987–4003. doi: 10.1128/IAI.72.7.3987-4003.2004

21. Diao J, Bouwman C, Yan D, Kang J, Katakam AK, Liu P, Pantua H, Abbas AR, Nickerson NN, Austin C, Reichelt M, Sandoval W, Xu M, Whitfield C, Kapadia SB. 2017. Peptidoglycan association of murein lipoprotein is required for KpsD-dependent group 2 capsular polysaccharide expression and serum resistance in a uropathogenic *Escherichia coli* isolate. mBio 8(3):e00603–17. doi: 10.1128/mBio.00603-17

22. Asmar AT, Collet JF. 2018. Lpp, the Braun lipoprotein, turns 50-major achievements and remaining issues. FEMS Microbiol Lett 365. doi.org/10.1093/femsle/fny199

23. Mizuno T, Kageyama M. 1979. Isolation of characterization of a major outer membrane protein of *Pseudomonas aeruginosa*. Evidence for the occurrence of a lipoprotein. J Biochem 85:115–22. doi: 10.1093/oxfordjournals.jbchem.a132300

24. Godessart P, Lannoy A, Dieu M, Van der Verren SE, Soumillion P, Collet JF, Remaut H, Renard P, De Bolle X. 2020. β-Barrels covalently link peptidoglycan and the outer membrane in the α-proteobacterium Brucella abortus. Nat Microbiol. 6,27–33(2021). doi.org/10.1038/s41564-020-00799-3

25. Sandoz KM, Moore RA, Beare PA, Patel AV, Smith RE, Bern M, Hwang H, Cooper CJ, Priola SA, Parks JM, Gumbart JC, Mesnage S, Heinzen RA. 2020. β-Barrel proteins tether the outer membrane in many Gram-negative bacteria. Nat Microbiol. 6, 19–26(2021). doi:10.1038/s41564-020-00798-4.

26. Schneewind O, Mihaylova-Petkov D, Model P. 1993. Cell wall sorting signals in surface proteins of gram-positive bacteria. EMBO J. 12:4803–4811. doi.org/10.1002/j.1460-2075.1993.tb06169.x

27. LeMieux J, Woody S, Camilli A. 2008. Roles of the sortases of *Streptococcus pneumoniae* in assembly of the RlrA pilus. J Bacteriol 190:6002–13. doi: 10.1128/JB.00379-08

28. Schneewind O, Missiakas D. 2019. Sortases, surface proteins, and their roles in *Staphylococcus aureus* disease and vaccine development. Microbiol Spectr 7(1):PSIB-0004-2018. doi: 10.1128/microbiolspec.PSIB-0004-2018

29. Moré N, Martorana AM, Biboy J, Otten C, Winkle M, Serrano CKG, Monton Silva A, Atkinson L, Yau H, Breukink E, den Blaauwen T, Vollmer W, Polissi A. 2019. Peptidoglycan remodeling enables *Escherichia coli* to survive severe outer membrane assembly defect. mBio 10(1):e02729–18. doi: 10.1128/mBio.02729-18

30. Guest RL, Same Guerra D, Wissler M, Grimm J, Silhavy TJ. 2020. YejM modulates activity of the YciM/FtsH protease complex to prevent lethal accumulation of lipopolysaccharide. mBio 11(2):e00598–20. doi: 10.1128/mBio.00598-20

31. Nicolaes V, El Hajjaji H, Davis RM, Van der Henst C, Depuydt M, Leverrier P, Aertsen A, Haufroid V, Ollagnier de Choudens S, De Bolle X, Ruiz N, Collet JF. 2014. Insights into the function of YciM, a heat shock membrane protein required to maintain envelope integrity in *Escherichia coli*. J Bacteriol 196:300–9. doi: 10.1128/JB.00921-13.

31b. Li G, Hamamoto K, Kitakawa M. 2012. Inner membrane protein YhcB interacts with RodZ involved in cell shape maintenance in *Escherichia coli*. ISRN Molecular Biology 304021. doi:10.5402/2012/304021

32. Monton Silva A, Otten C, Biboy J, Breukink E, VanNieuwenhze M, Vollmer W, den Blaauwen T. 2018. The fluorescent D-amino acid NADA as a tool to study the conditional activity of transpeptidases in *Escherichia coli*. Front Microbiol 9:2101. doi: 10.3389/fmicb.2018.02101

33. Peters K, Pazos M, Edoo Z, Hugonnet JE, Martorana AM, Polissi A, VanNieuwenhze MS, Arthur M, Vollmer W. 2018. Copper inhibits peptidoglycan LD-transpeptidases suppressing β-lactam resistance due to bypass of penicillin-binding proteins. Proc Natl Acad Sci U S A 115:10786–10791. doi: 10.1073/pnas.

34. Nichols RJ, Sen S, Choo YJ, Beltrao P, Zietek M, Chaba R, Lee S, Kazmierczak KM, Lee KJ, Wong A, Shales M, Lovett S, Winkler ME, Krogan NJ, Typas A, Gross CA. 2011. Phenotypic landscape of a bacterial cell. Cell 144:143–56. doi: 10.1016/j.cell.2010.11.052

35. Spratt BG. 1975. Distinct penicillin binding proteins involved in the division, elongation, and shape of *Escherichia coli* K12. Proc Natl Acad Sci U S A 72:2999–3003. doi: 10.1073/pnas.72.8.2999

36. Bean GJ, Flickinger ST, Westler WM, McCully ME, Sept D, Weibel DB, Amann KJ. 2009. A22 disrupts the bacterial actin cytoskeleton by directly binding and inducing a low-affinity state in MreB. Biochemistry 48:4852–7. doi: 10.1021/bi900014d

37. Lai GC, Cho H, Bernhardt TG. 2017. The mecillinam resistome reveals a role for peptidoglycan endopeptidases in stimulating cell wall synthesis in *Escherichia coli*. PLoS Genet 13:e1006934. doi: 10.1371/journal.pgen.1006934

38. Mendler K, Chen H, Parks DH, Lobb B, Hug LA, Doxey AC. 2019. AnnoTree: visualization and exploration of a functionally annotated microbial tree of life. Nucleic Acids Res 47:4442–4448. doi.org/10.1093/nar/gkz246

39. Rawlings ND, Barrett AJ, Thomas PD, Huang X, Bateman A, Finn RD. 2018. The MEROPS database of proteolytic enzymes, their substrates and inhibitors in 2017 and a comparison with peptidases in the PANTHER database. Nucleic Acids Res 46:D624–D632. doi.org/10.1093/nar/gkx1134

40. Kim HS, Im HN, An DR, Yoon JY, Jang JY, Mobashery S, Hesek D, Lee M, Yoo J, Cui M, Choi S, Kim C, Lee NK, Kim SJ, Kim JY, Bang G, Han BW, Lee BI, Yoon HJ, Suh SW. 2015. The cell shape-determining Csd6 protein from *Helicobacter pylori* constitutes a new family of L,D-carboxypeptidase. J Biol Chem 290:25103–17. doi: 10.1074/jbc.M115.658781

41. Frirdich E, Vermeulen J, Biboy J, Soares F, Taveirne ME, Johnson JG, DiRita VJ, Girardin SE, Vollmer W, Gaynor EC. 2014. Peptidoglycan LD-carboxypeptidase Pgp2 influences *Campylobacter jejuni* helical cell shape and pathogenic properties and provides the substrate for the DL-carboxypeptidase Pgp1. J Biol Chem 289:8007–18. doi: 10.1074/jbc.M113.491829

42. Ellis TN, Kuehn MJ. 2010. Virulence and immunomodulatory roles of bacterial outer membrane vesicles. Microbiol Mol Biol Rev 74:81–94. doi: 10.1128/MMBR.00031-09

42b. Sheikh J, Hicks S, Dall’Agnol M, Phillips AD, Nataro JP. 2001. Roles for Fis and YafK in biofilm formation by enteroaggregative *Escherichia coli*. Mol Microbiol 41(5):983–97. doi: 10.1046/j.1365-2958.2001.02512.x

43. Klein G, Kobylak N, Lindner B, Stupak A, Raina S. 2014. Assembly of lipopolysaccharide in *Escherichia coli* requires the essential LapB heat shock protein. J Biol Chem 289:14829–53. doi: 10.1074/jbc.M113.539494

44. Mahalakshmi S, Sunayana MR, SaiSree L, Reddy M. 2014. *yciM* is an essential gene required for regulation of lipopolysaccharide synthesis in *Escherichia coli*. Mol Microbiol 91:145–57. doi: 10.1111/mmi.12452

45. Cava F, de Pedro MA, Lam H, Davis BM, Waldor MK. 2011. Distinct pathways for modification of the bacterial cell wall by non-canonical D-amino acids. EMBO J 30:3442–53. doi: 10.1038/emboj.2011.246

46. Le NH, Peters K, Espaillat A, Sheldon JR, Gray J, Di Venanzio G, Lopez J, Djahanschiri B, Mueller EA, Hennon SW, Levin PA, Ebersberger I, Skaar EP, Cava F, Vollmer W, Feldman MF. 2020. Peptidoglycan editing provides immunity to *Acinetobacter baumannii* during bacterial warfare. Sci Adv 6:eabb5614. doi: 10.1126/sciadv.abb5614

47. Alvarez L, Aliashkevich A, de Pedro MA, Cava F. 2018. Bacterial secretion of D-arginine controls environmental microbial biodiversity. ISME J 12:438–450. doi: 10.1038/ismej.2017.176

48. Baba T, Ara T, Hasegawa M, Takai Y, Okumura Y, Baba M, Datsenko KA, Tomita M, Wanner BL, Mori H. 2006. Construction of *Escherichia coli* K-12 in-frame, single-gene knockout mutants: the Keio collection. Mol Syst Biol 2:2006 0008. doi: 10.1038/msb4100050

49. Silhavy TJ, Berman ML, Enquist LW. 1984. Experiments with gene fusions. CSH ColdSpringHarborLaboratory NY, Editor. ISBN-10: 0879691638

50. Datsenko KA, Wanner BL. 2000. One-step inactivation of chromosomal genes in *Escherichia coli* K-12 using PCR products. Proc Natl Acad Sci U S A 97:6640–5. doi.org/10.1073/pnas.120163297

51. Jeong JY, Yim HS, Ryu JY, Lee HS, Lee JH, Seen DS, Kang SG. 2012. One-step sequence- and ligation-independent cloning as a rapid and versatile cloning method for functional genomics studies. Appl Environ Microbiol 78:5440–3. doi: 10.1128/AEM.00844-12

52. Wurch T, Lestienne F, Pauwels PJ. 1998. A modified overlap extension PCR method to create chimeric genes in the absence of restriction enzymes. Biotechnology Techniques 12, 653–657. doi:10.1023/A:1008848517221:653-657.

53. Studier FW. 2005. Protein production by auto-induction in high density shaking cultures. Protein Expr Purif 41:207–34. doi.org/10.1016/j.pep.2005.01.016

54. . Tartoff, K.D. and Hobbs, C.A. 1987. Improved media for growing plasmid and cosmid clones. Bethesda Res Lab Focus 9.

55. Hugonnet JE, Mengin-Lecreulx D, Monton A, den Blaauwen T, Carbonnelle E, Veckerle C, Brun YV, van Nieuwenhze M, Bouchier C, Tu K, Rice LB, Arthur M. 2016. Factors essential for L,D-transpeptidase-mediated peptidoglycan cross-linking and β-lactam resistance in *Escherichia coli*. Elife 5:e19469. doi: 10.7554/eLife.19469

56. Bui NK, Gray J, Schwarz H, Schumann P, Blanot D, Vollmer W. 2009. The peptidoglycan sacculus of *Myxococcus xanthus* has unusual structural features and is degraded during glycerol-induced myxospore development. J Bacteriol 191:494–505. doi: 10.1128/JB.00608-08

57. Camacho C, Coulouris G, Avagyan V, Ma N, Papadopoulos J, Bealer K, Madden TL. 2009. BLAST+: architecture and applications. BMC Bioinformatics 10:421. doi.org/10.1186/1471-2105-10-421

58. Letunic I, Bork P. 2019. Interactive Tree Of Life (iTOL) v4: recent updates and new developments. Nucleic Acids Res 47:W256–W259. doi: 10.1093/nar/gkz239

59. Sievers F, Wilm A, Dineen D, Gibson TJ, Karplus K, Li W, Lopez R, McWilliam H, Remmert M, Soding J, Thompson JD, Higgins DG. 2011. Fast, scalable generation of high-quality protein multiple sequence alignments using Clustal Omega. Mol Syst Biol 7:539. DOI: 10.1038/msb.2011.75

60. Wheeler TJ, Clements J, Finn RD. 2014. Skylign: a tool for creating informative, interactive logos representing sequence alignments and profile hidden Markov models. BMC Bioinformatics 15:7. doi.org/10.1186/1471-2105-15-7

61. Tareen A, Kinney JB. 2020. Logomaker: beautiful sequence logos in Python. Bioinformatics 36:2272–2274. doi.org/10.1093/bioinformatics/btz921

62. Goodall ECA, Robinson A, Johnston IG, Jabbari S, Turner KA, Cunningham AF, Lund PA, Cole JA, Henderson IR. 2018. The essential genome of *Escherichia coli* K-12. mBio 9(1):e02096–17. doi: 10.1128/mBio.02096-17

63. Pearson WR, Wood T, Zhang Z, Miller W. 1997. Comparison of DNA sequences with protein sequences. Genomics 46:24–36. doi: 10.1006/geno.1997.4995

64. Barquist L, Mayho M, Cummins C, Cain AK, Boinett CJ, Page AJ, Langridge GC, Quail MA, Keane JA, Parkhill J. 2016. The TraDIS toolkit: sequencing and analysis for dense transposon mutant libraries. Bioinformatics 32:1109–11. doi: 10.1093/bioinformatics/btw022

